# Genomic characterization and therapeutic potential of five broad-spectrum lytic bacteriophages against multidrug-resistant avian pathogenic *Escherichia coli* (APEC)

**DOI:** 10.64898/2026.05.21.727054

**Authors:** Tushar Midha, Vishakha, Somesh Baranwal

## Abstract

Colibacillosis, caused by Avian Pathogenic *Escherichia coli* (APEC), result in substantial economic losses in global poultry production. The emergence of multidrug-resistant (MDR) APEC poses zoonotic risks through horizontal transfer of antimicrobial resistance (AMR) genes. Bacteriophage therapy emerges as a safe alternative to antibiotherapy; however, comprehensive characterization of phages targeting MDR-APEC from diverse geographical regions remains limited.

We isolated five lytic bacteriophages from poultry fecal samples collected from five Indian states and characterized them through morphological analysis, physiological stability testing, whole-genome sequencing, and *in vivo* efficacy assessment. Host range was determined against APEC isolates, and therapeutic potential was validated in *Galleria mellonella* infection model.

All five phages showed Myovirus-like morphology and stability across physiologically relevant temperatures (up to 55-70°C) and pH conditions (3-11). Their genome size ranges from 170 to 356 kb, belonging to three distinct genera; Dhakavirus, Gaprivervirus, and Asteriusvirus. Genomic analysis confirmed absence of antimicrobial resistance, virulence, toxin, or lysogeny genes. 51 APEC strains were isolated, of which 23 (45.1%) were MDR. Individual phages lysed 37-51% of tested APEC and 17-39% of MDR strains. Three *Escherichia* phages (fBSZT1, fUAMT1, fPKPT2) significantly improved larval survival to 60-80% at MOI 10 in *G. mellonella* infection models compared to untreated controls.

This study establishes a well-characterized phage bank targeting MDR-APEC strains, providing foundation for developing phage-based interventions to reduce antibiotic dependency and mitigate AMR transmission risks under One Health framework.

## 1. Introduction

Avian Pathogenic *E. coli* (APEC), a subset of Extra-intestinal Pathogenic *E. coli* (ExPEC), possesses virulence factors that enable it to cause systemic diseases such as colibacillosis, despite its potential commensal presence in the gut of some birds (1). APEC infections lead to high morbidity and mortality, reducing egg and meat production, causing significant economic losses to the global food and poultry industries (2).

In South Asian countries, indiscriminate use of antibiotics in broiler chickens as feed additives to promote growth and as a prophylactic agent to prevent infections puts selective pressure for the emergence of antibiotic-resistant bacteria (3). Most *E. coli* strains isolated from poultry exhibit multidrug resistance (MDR) to nearly all antibiotic classes, with only carbapenem, streptomycin, and tetracycline retaining marginal activity, making most conventional treatments ineffective (2). In India, a high resistance level is also observed against these antibiotics (4). The World Health Organization (WHO) has listed third-generation cephalosporin and carbapenem-resistant *E. coli* as critical priority pathogens (5). Further, commensal *E. coli* in the avian gut may act as a reservoir for antimicrobial resistance (AMR) genes, with potential to become pathogenic under stress (6). This poses a risk for zoonotic transfer of AMR, threatening animal and human health (2, 7). Antibiotic residues in poultry products and waste contaminate the environment (8, 9).

With the decline in new antibiotic development and the emergence of MDR bacteria, the poultry industry urgently requires alternative strategies to manage APEC infections (10). Currently explored interventions include large-scale vaccination, surface irradiation, strict biosecurity measures (including hygiene protocols, flock management, and environmental disinfection), and engineering chicken breeds with increased innate immune response to control vertical transmission of APEC (2, 11). Among these approaches, bacteriophage therapy has emerged as a promising alternative that can prevent infections and the horizontal transfer of AMR genes from animals to humans. Bacteriophages (phages), the viruses that infect bacteria, provide several advantages over conventional antibiotics (12). Phages have high host specificity, which minimizes disruption of natural microbiota, self-replication ability at infection sites, and a low likelihood of side effects. However, their narrow host range and potential resistance development by the bacteria make their therapeutic use challenging (13). Therefore, complete phenotypic and genotypic characterization and determination of host range become necessary for developing effective and safe phage-based therapies.

The U.S. Food and Drug Administration (FDA) has considered certain phages as Generally Recognized As Safe (GRAS) for food applications. Phages have been reported to control APEC both *in vitro* and *in vivo* (14, 15). Notably, a cocktail of four bacteriophages (FÓRMIDA) has shown promising *in vitro* results in controlling APEC in Brazil (1). However, despite over 15 ongoing clinical trials globally, no *E. coli* phage has received FDA approval for therapeutic use in the veterinary sector, although phage products like EcoShield™ PX have been approved for food safety applications (16). In India, comprehensive studies characterizing APEC prevalence, antibiotic resistance patterns, and phage susceptibility remain limited, creating a critical knowledge gap for the development of region-specific therapeutic solutions.

We report the morphological, bio-physicochemical, and genomic characteristics of five phages with therapeutic potential against MDR-APEC prevalent in Indian poultry. We validate the *in-vivo* efficacy of these phages using the *Galleria mellonella* infection model. This work addresses a critical need for alternative antimicrobial strategies in Indian poultry production and contributes to One Health framework by providing characterized phage resources that may reduce antibiotic consumption, control resistance development, prevent zoonotic AMR transfer, and minimize environmental contamination.

## 2. Results

### 2.1. Isolation and morphological characterization of bacteriophages

We isolated five lytic bacteriophages against host *E. coli* O135 from poultry faecal samples collected from four Indian states and a Union territory **(Supplementary Table S1)**. The phages were named as *Escherichia* phage fRPOT1 (fRPOT1), *Escherichia* phage fBSZT1 (fBSZT1), *Escherichia* phage fDMYT1 (fDMYT1), *Escherichia* phage fUAMT1 (fUAMT1), and *Escherichia* phage fPKPT2 (fPKPT2). All five phages showed a clear lysis zone (plaque) on host *E. coli* plates **(Fig.1)**. The phage fDMYT1 showed large plaques of 2.14 ± 0.13 mm **(Fig.1C)**. The diameters of plaques formed by phage fRPOT1, fBSZT1, fUAMT1, and fPKPT2 were 0.37 ± 0.03 mm, 0.22 ± 0.03 mm, 0.88 ± 0.1 mm, and 0.87 ± 0.08 mm, respectively **(Supplementary Table S1).** The phage titers were determined and observed to be (5.33 ± 0.98) Χ10^9^, (1.33 ± 0.11) Χ10^11^, (8.73 ± 0.46) Χ10^9^, (1.52 ± 0.91) Χ10^8^, (2.21 ± 0.192) Χ10^7^ PFU/mL for phage fRPOT1, fBSZT1, fDMYT1, fUAMT1, and fPKPT2, respectively. Further, transmission electron microscopy (TEM) results showed that these phages belong to Myoviruses with head-tail geometry **(Fig.1).** Similar to the plaque morphology, fDMYT1 showed a large head diagonal length of 107.62 ± 3.47 nm. In contrast, its tail length, i.e., 123.80 ± 3.96 nm, is comparable to other phages **(Fig.1C).** The tail length of phage fRPOT1, fBSZT1, fUAMT1, and fPKPT2 were 117.42 ± 2.26 nm, 116.23 ± 1.29 nm, 129.14 ± 8.25 nm, and 131.03 ± 1.05 nm while their head size is 75.46 ± 1.83 nm, 102.67 ± 1.07 nm, 92.18 ± 0.59 nm, and 96.46 ± 1.90 nm, respectively **(Supplementary Table S1).**

**Fig. 1:**
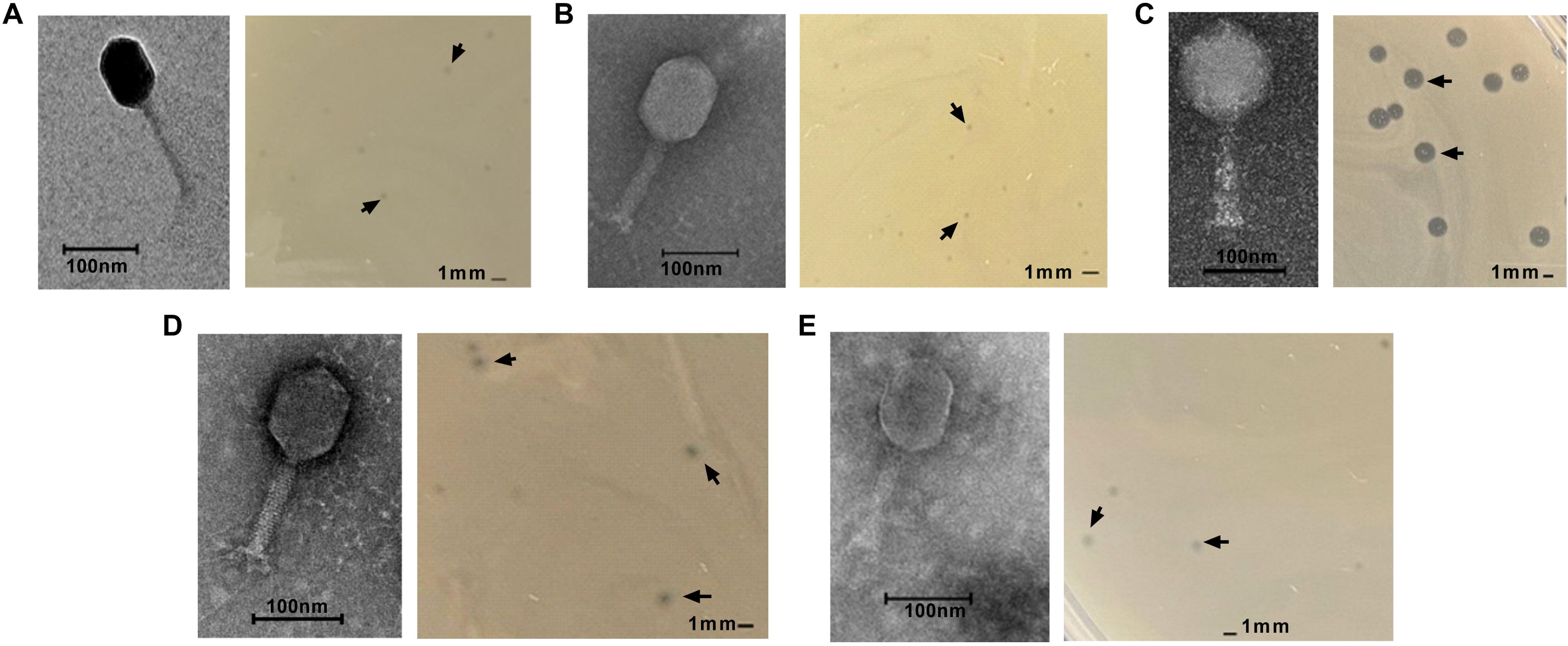
Morphological characteristics of five isolated *Escherichia* bacteriophages. Double layer agar overlay method was performed with 5 newly isolated bacteriophages, and bacterial lysis zones (plaques) were observed (indicated with black arrows). Images were taken with a 1mm scale bar. Transmission electron microscopy (TEM) revealed that all 5 phages that are **(A)** fRPOT1, **(B)** fBSZT1, **(C)** fDMYT1, **(D)** fUAMT1, and **(E)** fPKPT2 are Myovirus type. The TEM image after uranyl acetate staining is shown for each phage with a 100nm scale bar.

### 2.2. Bacteriophages show activity under extreme physiological conditions and long-term stability in the liquid state

Phages should be stable and active under physiological conditions to be used for veterinary applications, so we measured the phage activity after exposing them to extreme physiological pH and temperatures. All five phages remained stable at temperatures up to 55°C **(Fig.2A),** and phages fUAMT1 and fPKPT2 were stable up to 70°C. In addition, all phages showed stability at pH levels ranging from 3 to 11, except phage fDMYT1, which was stable at pH 4-11 **(Fig.2B).** Importantly, the phages maintained their titers up to 18 months when stored in SM buffer at 4°C **(Supplementary Fig. S1)**. After which 1-2 log reduction in plaque forming units, PFU/mL was observed. Similarly, when stored in 15% glycerol, the phages were stable for 18 months at -20°C and for 12 months at -80°C. Only phage fPKPT2 showed lower long-term stability under all conditions. It was stable for 12 months, and then a 3-4 log reduction in PFU/mL was observed **(Supplementary Fig. S1 E)**. We next determined phage adsorption rate using host *E. coli* O135 **(Fig.2C).** Phage fBSZT1 was 50% adsorbed by the host within 2 minutes of infection, while the other took around 4-8 min. This was also observed with the latent period and burst size as the phage fBSZT1 had the shortest latent period of 10 minutes and the highest burst size of 915.40 ± 252.22 phages/cell. While all other phages had a 20-min latent period and the burst size ranges from 62.95 ± 17.42, 556.02 ± 71.40, 92.23 ± 51.89, and 414.54 ± 183.23 phages/cell for phage fRPOT1, fDMYT1, fUAMT1, and fPKPT2, respectively **(Fig.2D).** In summary, our results support suitability of phages for veterinary applications due to their short latent period, rapid burst time and long-term activity in the liquid state.

**Fig. 2:**
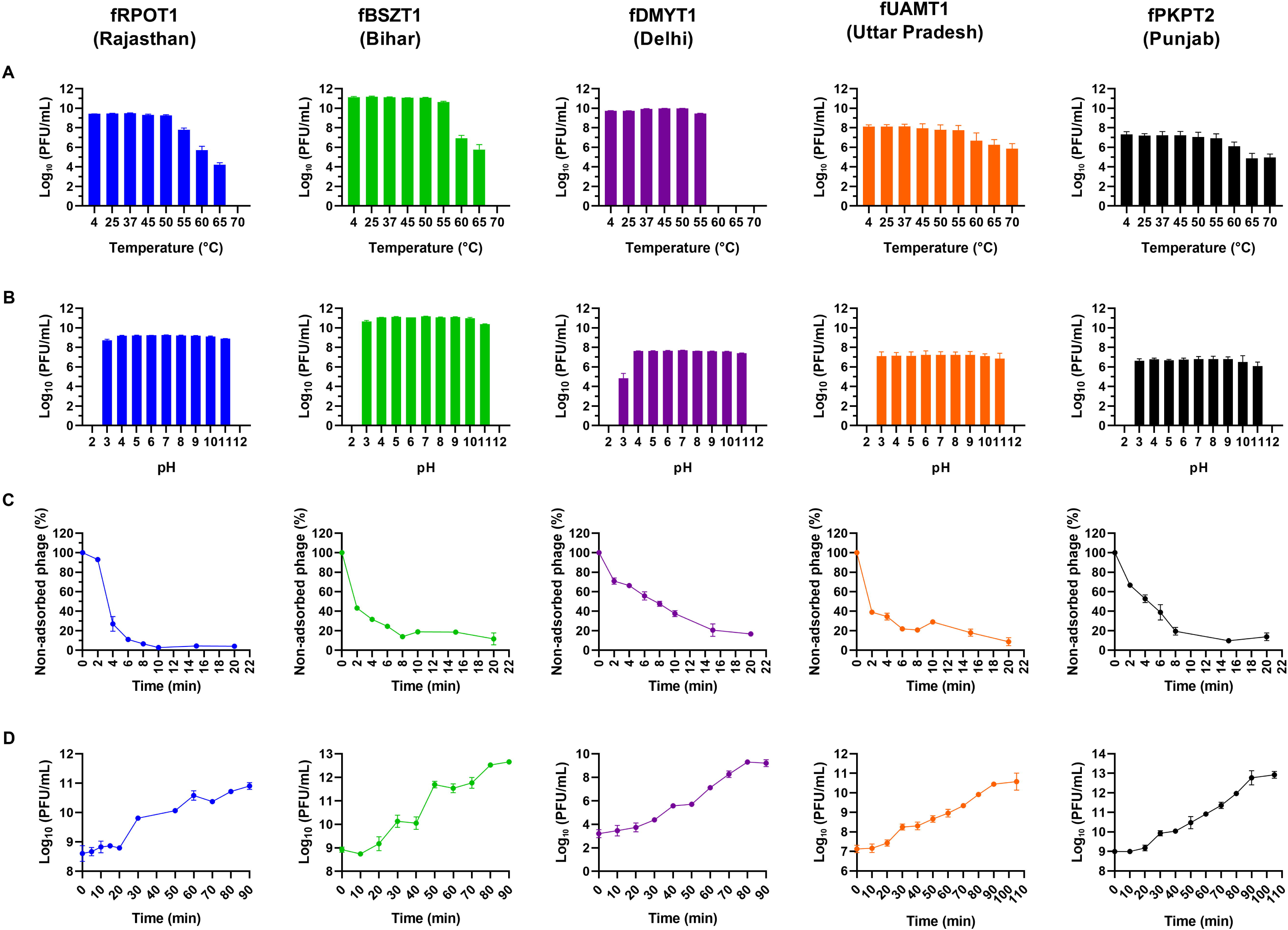
Physiological characteristics of isolated *Escherichia* phages. The physiological characteristics of 5 newly identified *Escherichia* phages are depicted here in 5 different columns, where the column heading shows the phage name and the Indian state from which the poultry fecal sample was collected for phage isolation. The row **(A)** represents the effect of different temperatures, while row **(B)** represents the effect of pH on phage stability. The plots in rows **(C)** and **(D)** present the number of unadsorbed phages with time and the phage one-step growth curve, respectively.

### 2.3. Disruption of bacterial cell surface proteins or carbohydrates leads to decreased phage adsorption

Recognition of phages with cell surfaces is crucial for successful lysis of gram-negative bacteria, so we determined how disruption of outer membrane proteins (OMPs) and carbohydrate by proteinase K and sodium periodate, respectively, affects the adsorption of phages. We observed a tenfold reduction of phage adsorption as shown by an increase in the number of unadsorbed phages after proteinase K treatment in all phages, indicating a potential role of OMPs in initiating the phage infection cycle **(Supplementary Fig. S2)**. However, since adsorption was not abolished entirely, incomplete enzyme access or a secondary receptor contribution cannot be excluded. For four of the five phages (fRPOT1, fBSZT1, fUAMT1, fPKPT2), sodium periodate treatment did not result in a significant reduction of phage adsorption, indicating that LPS is unlikely to be the primary receptor for these phages. fDMYT1, a member of the jumbo phage, adsorption was also affected by the disruption of carbohydrate component with sodium periodate **(Supplementary Fig. S2C)**.

### 2.4. Genomic studies reveal phage suitability for therapeutic applications

To confirm the genomic properties and rule out the existence of detrimental genetic material, we performed whole genome sequencing using Illumina platform (17). *De-novo* assembly and annotation revealed that phage fRPOT1, fBSZT1, fDMYT1, fUAMT1, and fPKPT2 have a genome size of 170501, 169720, 356482, 170021, 171101 bp having 279, 271, 588, 290, 283 coding regions (CDS) and G+C content of 39.55%, 39.65%, 35.60%, 40.38%, 40.26%, respectively **(Supplementary Table S1)**. All five phages were observed to have a conserved core replication machinery characteristic of T4-like Myoviruses, including a DNA-directed DNA polymerase, sliding clamp loader (gp44), DNA polymerase clamp, DNA primase, ATP-dependent helicases (gp41 and UvsW), recombination proteins (UvsX/UvsY homologues), and a complete nucleotide metabolism module comprising thymidylate synthase, thymidylate kinase, and dihydrofolate reductase. Structurally, all phages encode a complete baseplate assembly, tail sheath, tail tube, terminase large and small subunits, portal protein, major capsid protein, and a holin–endolysin–spanin lysis cassette. Despite these conserved genes, some key differences were observed among the phages. The jumbo phage fDMYT1 encodes a colanic acid degradation enzyme which is a capsule depolymerase. The *Escherichia* phages fBSZT1 and fUAMT1 have an anti-restriction nuclease which could counter host immune defences like restriction-modification systems, while phage fPKPT2 additionally encodes three homing endonucleases indicating a greater genomic mobility **(Supplementary Table S2)**. The phage genomic map was constructed using the Proksee CG view program, highlighting the identified protein functions and some major proteins **(Fig.3).** All phages code for 2 tRNAs, except phage fDMYT1 & fRPOT1, which encode 4 tRNAs. The phages lacked genes associated with AMR, virulence, toxicity, or lysogeny, suggesting their potential use as a therapeutic agent **(Supplementary Table S2)**. The phage’s lifestyle was confirmed to be completely lytic through the Life Cycle Classifier v1.6.0 accessed through PhageAI 1.0.2 **(Project accession PROJ-00232**) (18). These results suggest the lytic nature of all phages with no genes hindering their therapeutic use.

**Fig. 3:**
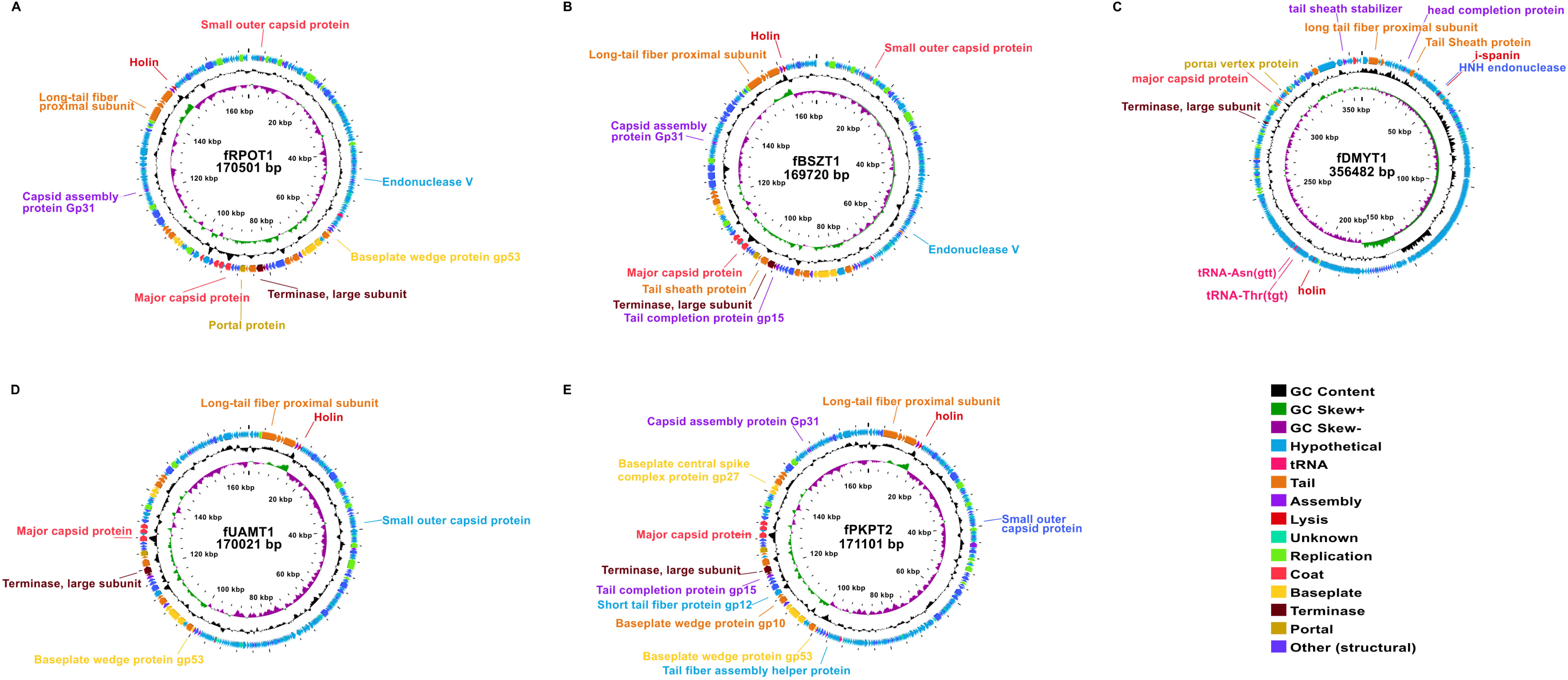
Genomic structure of *Escherichia* phages. The genomic organization of all *Escherichia* phages **(A)** fRPOT1, **(B)** fBSZT1, **(C)** fDMYT1, **(D)** fUAMT1, and **(E)** fPKPT2 is exhibited here. The genomic map was built with the Proksee CGview program, Map Builder 2.0.5. The inner layer represents the GC skew, going outward, the GC content and phage coding regions depicting their specific functions with different colors are shown here. Some phage core proteins are labelled in the genomic map, and the ruler stating phage length is also shown.

### 2.5. Phylogenetic and comparative analysis of *Escherichia* phages

To delineate the phylogenetic relationships of our phage, we constructed a viral circular proteomic tree of 5,637 phages **(Supplementary Table S3),** including our phages (highlighted with a red star), through ViPTree **(Fig.4)**. All 5 phages have a host group of Pseudomonadota. The phages fPKPT2 and fUAMT1 have a close relationship with each other compared to phages fRPOT1 and fBSZT1, which show similarity with other *Escherichia* phages. The genomic size of all phages in this clade ranges from 169 to 171.5 kbp, and they are members of Straboviridae family. The phage fDMYT1 shows similarity to *Escherichia* jumbo phages, PBECO4 and 121Q **(Fig.4)**. Finally, we confirm the assembly quality of our phages with closely related phages through the DigAlign 2.0 server **(Fig.4)**.

**Fig. 4:**
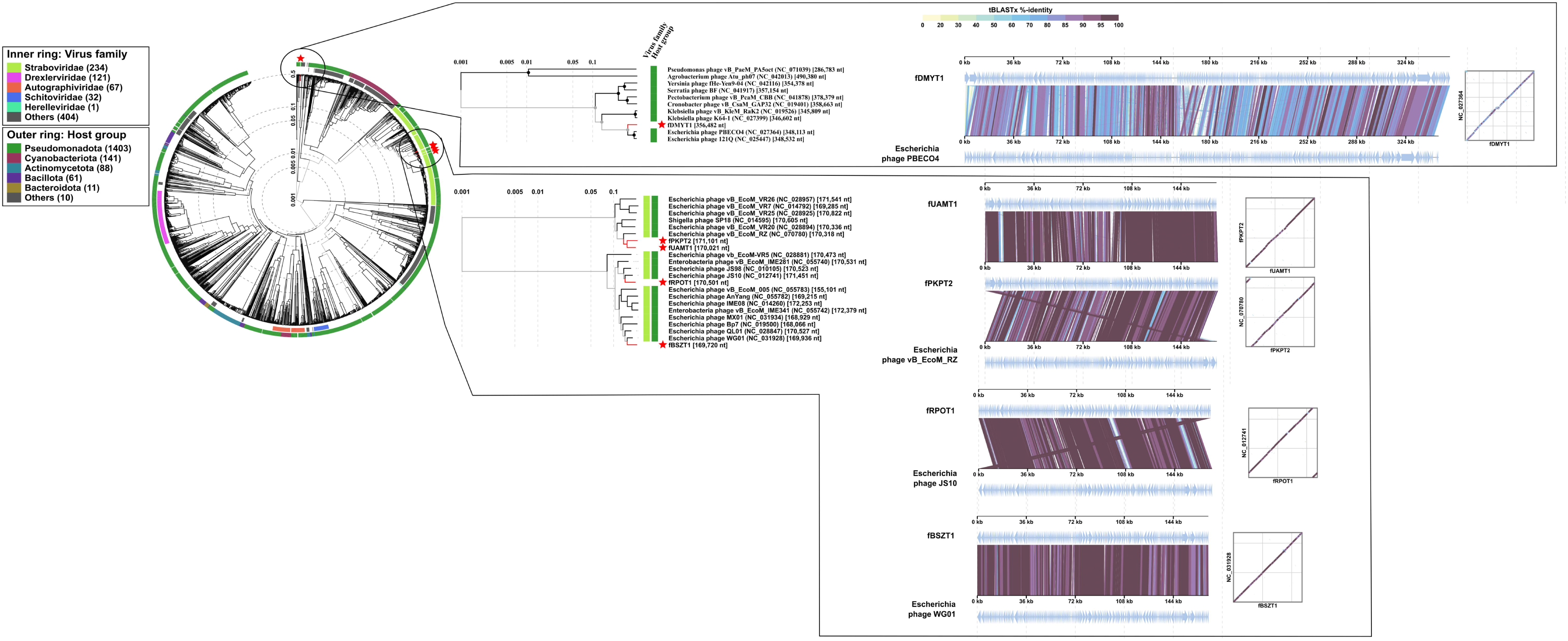
Phage proteomic tree and alignment. A circular phage proteomic tree is built with VipTree version 4.0 based on genome-wide sequence similarities computed through tBLASTx with reference viral genomes. The phage families and host groups are identified. Further, the rectangular tree shows the similarity observed with phages in the same taxa. The closely related phage genome for each of our 5 *Escherichia* phages was determined with *S_g_* score observed on proteomic analysis, and the alignment was built through DigALign version 2.0. The alignment of each phage with its closely related phage is displayed along with the dot plot.

We next aligned phage genomes with BLASTn, and sequences with >90% query coverage and percent identity were selected to determine the inter-genomic similarities with Virus Intergenomic Distance Calculator (VIRIDIC) **(Fig.5A)**. The phage fRPOT1 shows the highest similarity of 91.8% with Enterobacteria phage JS10 (NC_012741.1), phage fBSZT1 of 92.6% with *Escherichia* phage WG01 (KU_878968.1), phage fDMYT1 of 77.5% with *Escherichia* phage PBECO 4 (NC_027364.1), phage fUAMT1 of 96.4% with *Escherichia* phage pEC M719 6WT.2 (OQ_845958.1), and phage fPKPT2 of 90.2% with *Escherichia* phage EAWsc9 (PQ_341120.1) **(Supplementary Table S4)**. Further, the phage fRPOT1 and fBSZT1 show a sequence similarity of 85.3%, while phage fUAMT1 and fPKPT2 were more closely related with an inter-genomic similarity of 90.1%. Similar results were obtained upon performing genome-based phylogeny and classification through VICTOR **(Fig.5B)**. The Genome-BLAST Distance Phylogeny (GBDP) analysis of our phages with 49 closely related phages yields a tree of two genus clusters out of 50 species clusters **(Supplementary Table S5)**.

**Fig. 5:**
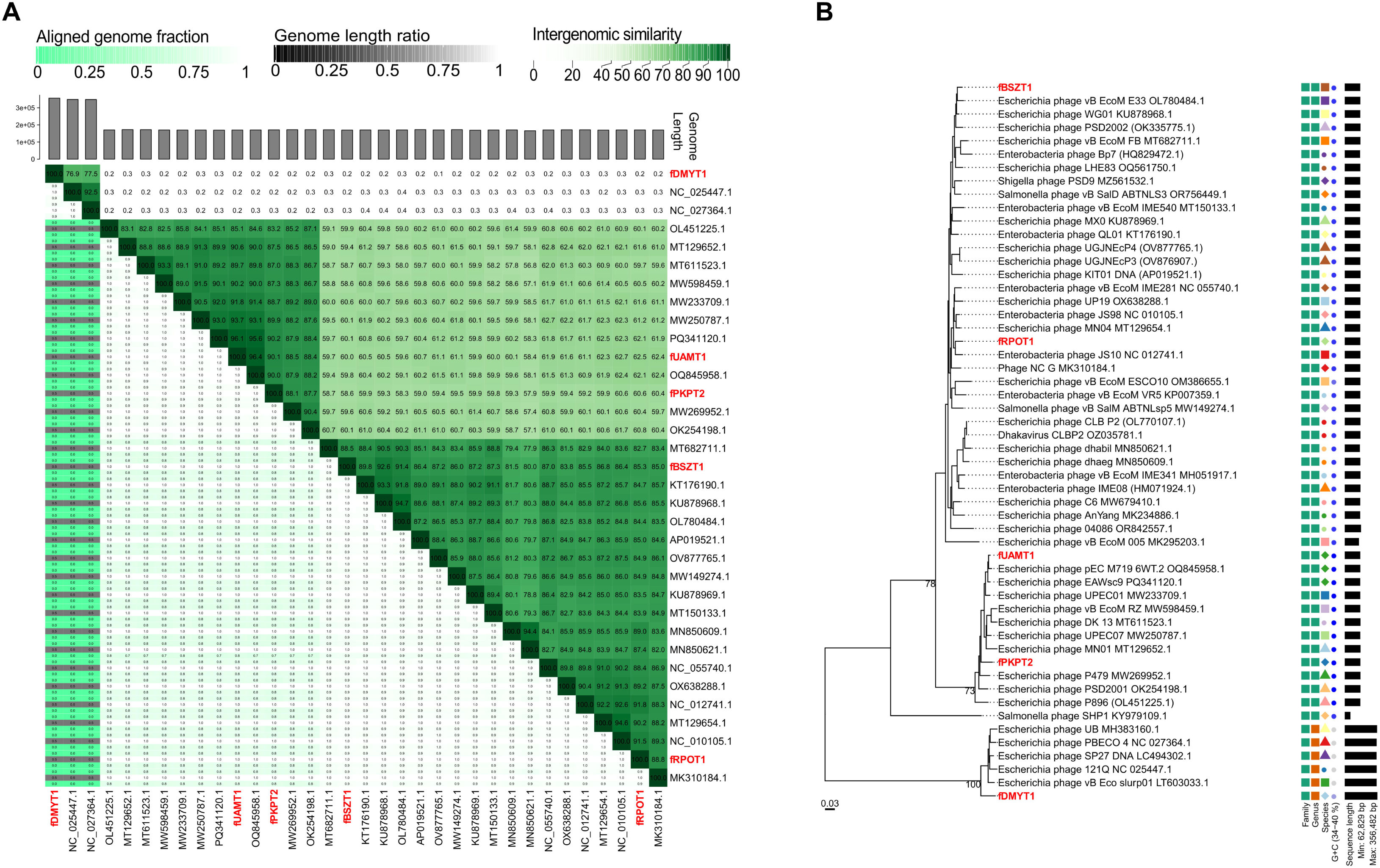
Intergenomic similarities and phylogeny of *Escherichia* phages. **(A)** The intergenomic similarity between the 5 isolated *Escherichia* phages and their closely related phages in the same taxa is shown, determined by BLASTn search through the NCBI database (>90% identity) and computed using Virus Intergenomic Distance Calculator (VIRIDIC). The intergenomic similarities (highlighted in dark green) are depicted. **(B)** The phylogenomic Genome-BLAST Distance Phylogeny (GBDP) trees inferred using the formulas D0 through VICTOR web service. The numbers above branches are GBDP pseudo-bootstrap support values from 100 replications. The branch lengths of the VICTOR tree is scaled in terms of the GBDP distance formula *d_0_*. The OPTSIL clustering yielded 50 species clusters and 2 clusters at the genus and 1 at the family level.

Though the identified most closely related phage for fRPOT1: NC_012741.1, and for fUAMT1:OQ_845958.1 were same by VIRIDIC **(Fig.5A)** and VICTOR **(Fig. 5B),** the two tools occasionally identified different closest relatives for the same phage; for example, the closest relative of fBSZT1 by VICTOR was OL780484.1, which was the 2^nd^ most closest by VIRIDIC. For fPKPT2 and fDMYT1, the differences observed were greater. This could be possible as VIRIDIC computes pair-wise nucleotide-level similarity, which is based on BLASTn result for whole genome (19), while VICTOR predicts the phylogeny by dividing high-scoring segment pairs with total genome lengths and is optimized against a large reference dataset of genome-sequenced taxa recognized by the International Committee on Taxonomy of Viruses (ICTV) (20).

Further, we generated a neighbour-joining tree using two protein orthologs, MCP and TerL, that were present in all phages. The tree produced for the TerL protein of all phages is given in **Fig.6A, B, C,** while for the MCP protein, the tree is shown in **Fig.6D, E**. The phages fRPOT1 and fBSZT1 were predicted to belong to the same genus by GBDP, which was confirmed as Dhakavirus by taxMyPhage 3.3.6 software analysis. The phages fUAMT1 and fPKPT2 belong to the genus Gaprivervirus, and are more closely related. They are predicted under the same taxa by VICTOR and has 99% branch support for phylogeny based on MCP **(Fig.6D)**. The *Escherichia* phage fDMYT1 belongs to the genus Asteriusvirus.

**Fig. 6:**
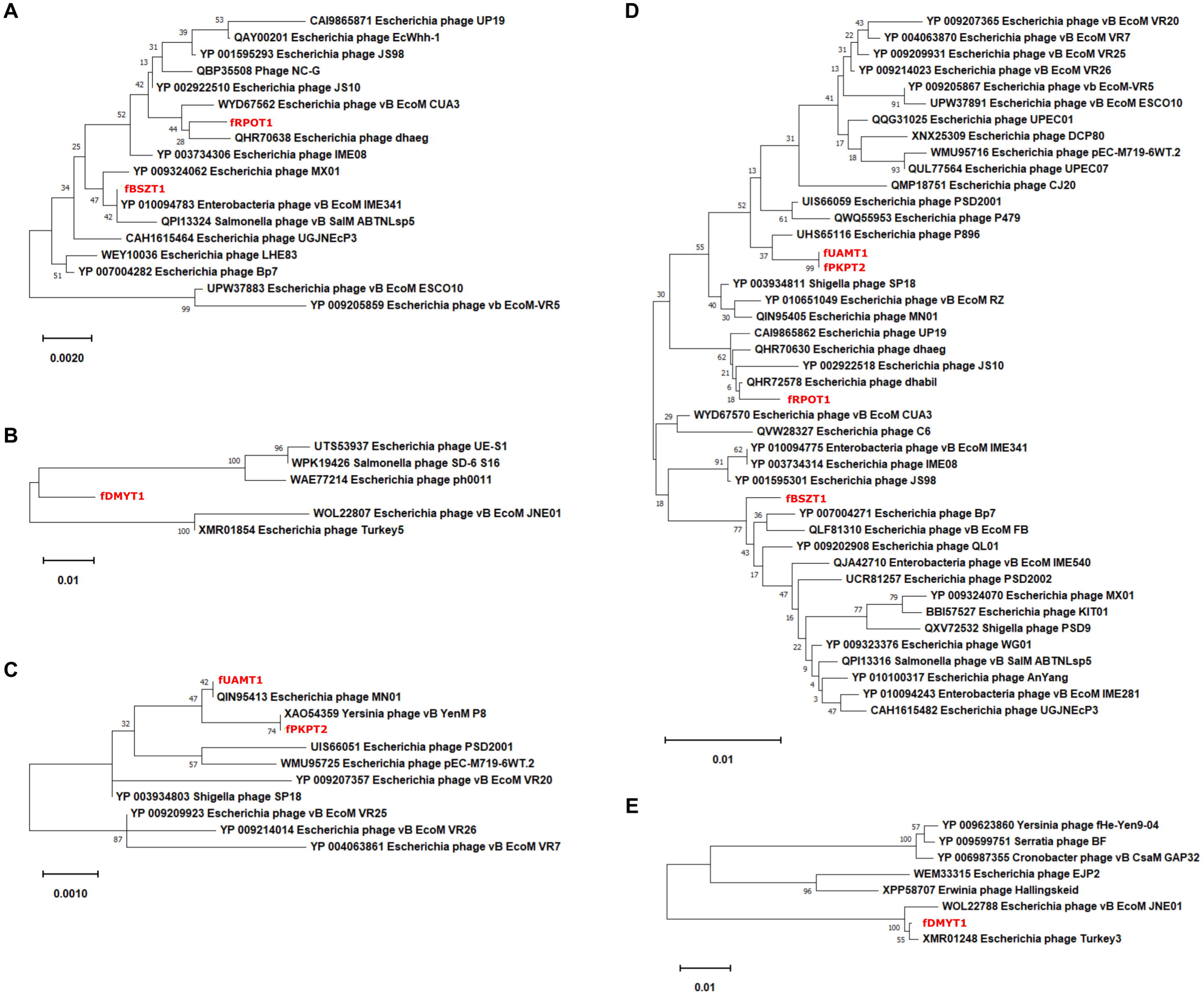
The phylogenetic relation and classification of phages based on two structural proteins: terminase large subunit (TerL) and major capsid protein (MCP). The amino acid sequence of TerL and MCP was aligned with BLASTp to the NCBI database, and closely related phages were identified (>90% identity). The protein sequence was aligned with muscle, and phyloanalysis was done with MEGA 11.0.13. **(A), (B)** and **(C)** represent the result obtained for the TerL protein for phage fRPOT1, fBSZT1, fDMYT1, fUAMT1, and fPKPT2, while **(D)** and **(E)** represent the phyloanalysis of the MCP protein of all 5 phages. Evolutionary history was inferred using the Neighbor-Joining method, and the percentage of replicate trees in which the associated taxa clustered together in the bootstrap test (1000 replicates) is shown next to the branches. The tree is drawn to scale, with branch lengths in the same units as those of the evolutionary distances used to infer the phylogenetic tree.

With intergenomic similarities from VIRIDIC, GBDP from VICTOR, and a phylogeny based on protein orthologs, we provide a more rigorous and multidimensional understanding of phage relatedness, suggesting that these five phages constitute distinct evolutionary lineages with some close relationship. Based on combined morphological, genomics, and phylogenetic evidences, we concluded that these lytic phages represent new *Escherichia* phages isolated from India.

### 2.6. Phages possess broad-spectrum activity against MDR APECs

To assess the lytic potential of our phages, we isolated *E. coli* strains from poultry faecal samples collected from infected chickens across five Indian states and one union territory **(Supplementary Table S6)**. The isolates were initially looked for presence of 3 APEC specific virulence genes, *hly*F, *omp*T, and *iro*N. 51 isolates were identified having all 3 genes. Further, they were looked for the presence of 2 additional APEC virulence genes, *iut*A and *iss* **(Supplementary Fig. S3)**. All 51 isolates had at least 4 of the 5 virulence factors and were considered APEC (21). APEC strains were serotyped at the **<u>N</u>**ational ***<u>S</u>****almonella* and ***<u>E</u>****scherichia* **<u>C</u>**entre (NSEC) at CRI, Kasauli. Surprisingly we identified 2 APEC isolates to be of serotype O156, which is usually associated with enterohemorrhagic *E. coli* (EHEC) (22). The presence of APEC virulence genes in *E. coli* O156 indicate a possible horizontal gene transfer. We measured the susceptibility of 51 isolates to 10 commonly used antibiotics representing 8 different drug classes using the Kirby-Bauer disk diffusion assay. The antibiotic resistance prevalence observed was: ciprofloxacin (CIP, 47.06%), colistin (CL, 47.05%) nitrofurantoin (NIT, 45.10%), tetracycline (TE, 41.17%), ceftazidime (CAZ, 29.41%), cefuroxime (CXM, 21.57%), amoxicillin/clavulanic acid (AMC, 21.56%), cotrimoxazole (COT, 19.61%), piperacillin/tazobactam (PIT, 9.8%), and ertapenem (ETP, 1.96%) **(Supplementary Fig. S4 A)**. Based on these results, we calculated the Multiple Antibiotic Resistance Index (MARI), which is defined as the ratio of resistant antibiotics to the total antibiotics tested. 28 strains exhibited MARI<0.3, while 23 showed MARI>0.3, which suggests they are resistant to over 3 antibiotics from different class out of the 10 antibiotics tested, making them MDR strains **(Supplementary Fig. S4 B)**. Given the prevalent use of nitrofurans in the poultry sector, we further tested 22 highly resistant strains against 8 additional antibiotics that belongs to fluoroquinolones and nitrofuran. Notably, the resistance prevalence to nitrofurans, furazolidone (FR), and nitrofurazone (NR) was universal, with no strain showing susceptibility to either antibiotic **(Supplementary Fig. S5)**.

Further, we performed a six-hour spot assay to check if phages can lyse APEC serotype **(Fig.7)**. Phages showed predominantly strong lytic activity, producing a clear zone of inhibition, though some moderate or slight lytic activity was also observed **(Supplementary Fig. S4 C)**. Out of 51 APEC strains, 37.25%, 45.10%, 47.06%, 50.98%, and 50.98% were susceptible to phage fDMYT1, fBSZT1, fUAMT1, fPKPT2, and fRPOT1 **(Supplementary Fig. S4 C)**. The strains belong to 17 different serotypes, and phages show activity against most of these serotypes except O169, O17, O20, and O9 **(Supplementary Fig. S4 D)**. Out of 23 MDR strains, 17.39%, 30.43%, 26.09%, 39.13%, and 34.78% were susceptible to phage fDMYT1, fBSZT1, fUAMT1, fPKPT2, and fRPOT1. The phages could lyse strains with all MARI values, with phage fPKPT2 being able to lyse strains having MARI of 0.7 and 0.9 **(Supplementary Fig. S4 B)**. *In-silico* Cytoscape PhageCocktail analysis shows that there are 18 bacterial strains that cannot be lysed by any of the bacteriophage, while the bacterial strains bPGOV1 and bPGOV2 can only be lysed by fPKPT2 and strain bDKRS2 is only susceptible to phage fUAMT1 **(Supplementary Fig. S6)**.

**Fig. 7:**
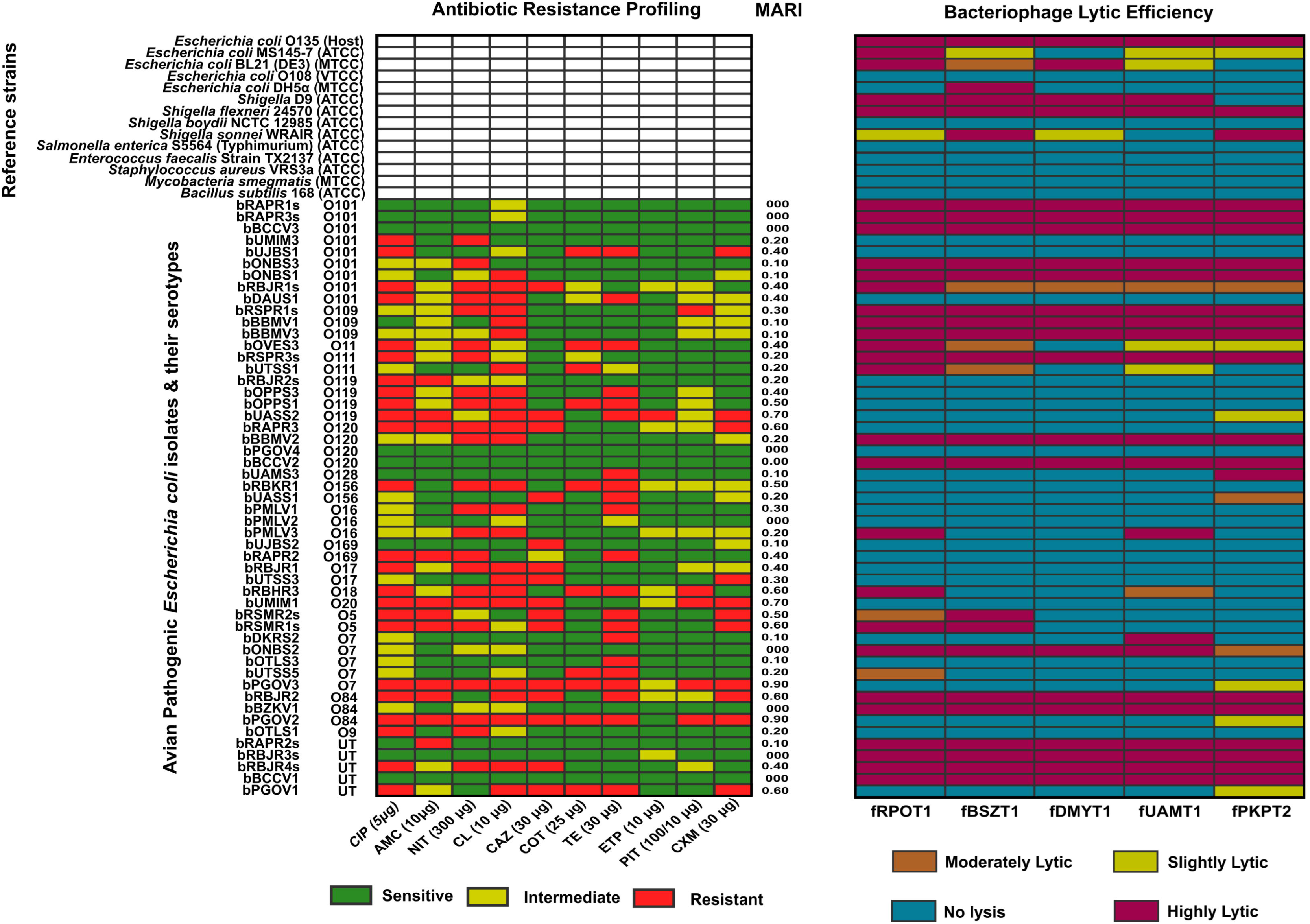
Bacteriophage host range against MDR-APEC. 51 Avian Pathogenic *E. coli* (APEC) were isolated, serotyped and their drug sensitivity profiling against 10 antibiotics belonging to different classes, fluoroquinolones (ciprofloxacin-CIP), penicillins with beta-lactamase inhibitor (amoxicillin/clavulanic acid-AMC, piperacillin/tazobactum-PIT), nitrofurans (nitrofurantoin-NIT), polymyxins (colistin-CL), cephalosporins (ceftazidime-CAZ, cefuroxime-CXM), sulfonamides (cotrimoxazole-COT), tetracyclins (tetracyclin-TE) and carbapenems (Ertapenem-ETP) is done. The Multiple Antibiotic Resistance Index (MARI) is calculated for each isolate. The effect of phage against each bacterial isolate was checked with a spot assay. The observed lysis zone is scored as complete lysis (purple), moderate lysis (Brown), turbid lysis (yellow), and no lysis (blue). The phage lysis against some reference strains is also determined.

For therapeutic development and veterinary applications, it is important to determine the host growth suppression in the liquid state, so we examined the growth kinetics of *E. coli* O135 host in presence of five phages (at multiplicity of infection (MOIs) of 0.1, 1, and 10). We observed a significant reduction in bacterial growth with all phages, even at an MOI of 0.1, compared to the SM buffer-treated bacterial host control over 48 h. **(Fig.8)**. However, there was an increase in bacterial growth observed after 8-12 h in all bacteriophages, probably due to the development of phage resistant mutants. Overall, these results suggest varying lytic ability of 5 phages from MDR-APEC strains isolated from several geographical locations in India, and highlight phage resistance as a consideration for any therapeutic application. Based on the bacterial susceptibility to antibiotics and phages, we measured their Pearson correlation **(Supplementary Table S7)**. A correlation between phage fRPOT1 and fBSZT1, and phage fUAMT1 and fPKPT2 activity is observed, which also corresponds to the correlation observed through inter-genomic similarities between these phages. Apart from these, CIP and TE showed a slight correlation with all phages, with TE showing the highest correlation with phage fDMYT1 activity. A negative correlation was observed between the activity of AMC and phage fRPOT1, fBSZT1, fUAMT1, and fPKPT2 against 51 APEC isolates. The phage fUAMT1 activity also shows a negative correlation with antibiotics CL and ETP. Two strains, bPGOV3 and bPGOV2, having serotypes O7 and O84, respectively, are only susceptible to antibiotic ETP and phage fPKPT2 **(Fig.7)**. These correlations may warrant future experimental investigation of phage-antibiotic combinations.

**Fig. 8:**
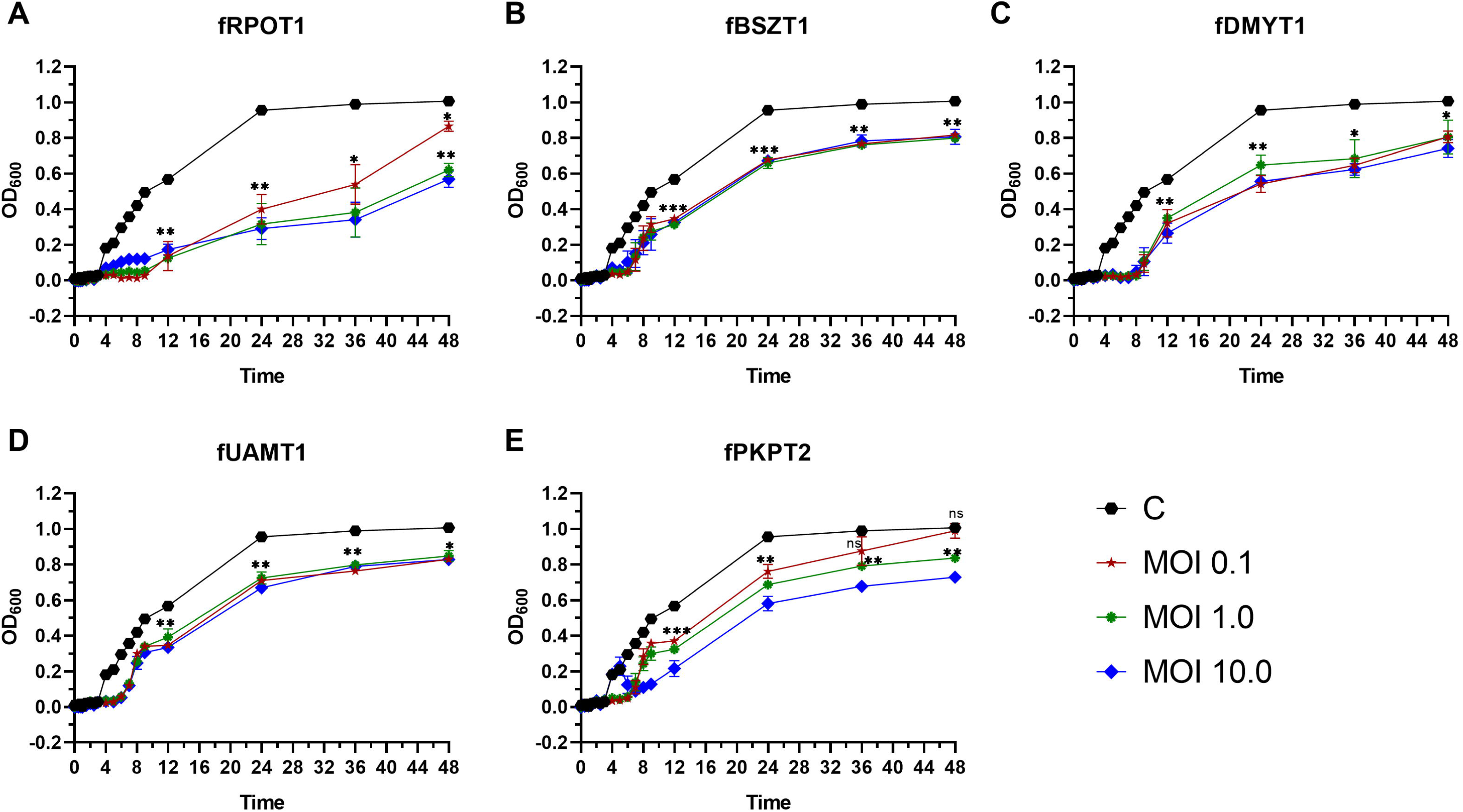
Lytic efficiency of *Escherichia* phages. A liquid suppression assay was performed to determine the inhibition of host *E. coli* with respect to time at different MOIs of 0.1, 1.0, and 10.0 of five isolated phages, namely **(A)** fRPOT1, **(B)** fBSZT1, **(C)** fDMYT1, **(D)** fUAMT1, and **(E)** fPKPT2. The absorbance at 600nm was measured at different time intervals. A significant reduction in bacterial growth was observed upon incubation at 37°C with phages post 6h for all phages up to 48h (*p<0.05, **p<0.01, ***p<0.001,****p<0.0001).

### 2.7. Phages reduce E. coli infection in G. mellonella in-vivo model

*G. mellonella* larvae phage infection model provides a simple *in-vivo* model to test the bacteriophage efficacy against bacterial pathogens. We first performed a dose-dependent assay to determine the appropriate bacterial count that causes 100% mortality in *G. mellonella* larval infection **(Supplementary Fig. S7)**. The larvae were infected with 10 microliters of *E. coli* (5X10^5^ CFU/ml) followed by phage injection at MOI 10 and MOI 1 after 1 h of post-infection (PI). At MOI 1, phage treatment did not significantly improve larval survival (data not shown), despite showing bacterial reduction *in vitro* (P<.00001). At 10 MOI, there was a log-fold decrease in bacterial count after treatment with phages fRPOT1, fBSZT1, and fDMYT1 as compared with the initial infected bacterial cell **(Supplementary Fig. S8)**, but the larvae’s survival was not significant with fRPOT1 and fDMYT1. The treatment with phage fBSZT1 at MOI of 10 led to 60% survival of larvae **(Fig.9)**. The phages fUAMT1 and fPKPT2 treatment showed a 1.5 and 2.25 log reduction in bacterial count, and 80% survivability of the larvae **(Fig.9).**

**Fig. 9:**
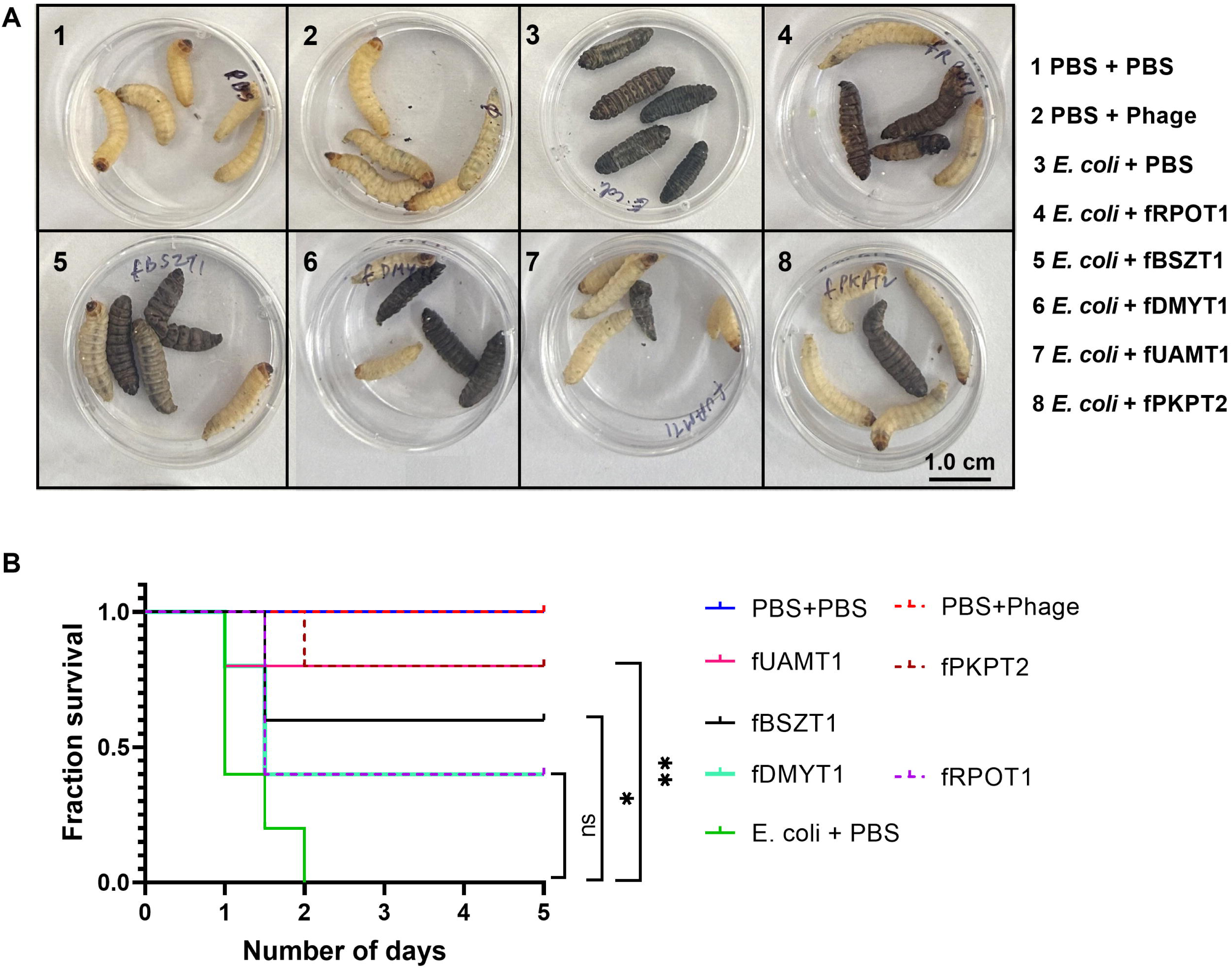
Efficacy of phages in treating *Escherichia coli* infection in *Galleria mellonella in vivo* model. The *G. mellonella* larvae were infected with *E. coli*, and 1 h post-infection (PI), a phage was injected at MOI 10 in larvae, and the survivability was observed for 5 days. **(A)** The images of larvae in each set are shown, depicting the melanised and dead larvae 5 days post-infection. **(B)** A Kaplan-Meier survival plot is shown with a log-rank test performed to determine significance, \*\**p*<0.001, \**p*<0.05. The larvae injected with phage fUAMT1 and fPKPT2 show the highest survivability (*p*<0.001).

## 3. Discussion

Because of a surge in demand for poultry products and the uncontrolled use of antibiotics, the veterinary sector poses a huge risk for the emergence of multidrug resistance among APEC. Bacteriophage-based formulations, already approved as GRAS category by FDA, have recently emerged as a promising alternative to minimise antibiotic dependency and protect the environment from antibiotic leaching in the agricultural, food safety, and veterinary sectors.

In this study, we isolated five APEC-specific phages from chicken faecal samples collected across multiple Indian states and characterized them comprehensively at phenotypic, genomic, and functional levels. These phages possess favourable properties for therapeutic development, such as broad stability across temperatures, pH ranges, and rapid adsorption, and their genomes lack any genes associated with lysogeny, virulence, toxicity, or antimicrobial resistance. Critically, phages showed *in vitro* efficacy against diverse MDR-APEC isolates and protected larvae in an *in vivo* infection model, validating their therapeutic potential. However, some phage resistance was observed in lysis kinetics and *in vivo* studies, which needs to be further studied, and a phage cocktail could be developed to overcome these limitations.

All five phages showed Myovirus-like morphology, and varied in lytic activity and physicochemical properties such as temperature and pH stability, adsorption rate, and burst size, showing differences in their growth dynamics and activity. They belong to three different genera; Dhakavirus, and Gaprivervirus within the Straboviridae family, and Asteriusvirus, reflecting the phylogenetic diversity among *Escherichia* phages in India. One of our phages, fDMYT1, is a jumbo phage with a 356 kb genome. Jumbo phage has a complex genomic structure with multiple paralogs of essential phage life cycle genes and genes with associated functions scattered around the genome (23). fDMYT1 possesses the same complexity. Phages may use multiple receptor binding proteins (RBPs), which could contribute to their broad host range (24). In contrast to four phages whose adsorption was affected by only bacterial surface protein disruption, fDMYT1 adsorption reduced on both host surface protein and carbohydrates disruption, emphasizing the possibility of a dual receptor. However, fDMYT1 showed narrow host range (37.25%). This suggests either the low prevalence of its specific receptors targets among Indian APEC strains or the presence of phage defence systems in non-susceptible strains.

The bacteriophage plaque size is dependent on multiple factors like latent period, adsorption rate, burst size, virion size, and diffusion rate through agar (25). Usually the phages with larger size slowly diffuse in the media and produces smaller plaques but the phage fDMYT1 had large plaque size despite having a relatively large head diameter and small burst size, while fBSZT1, which had the largest burst size, produced small plaques. The contradiction in fDMYT1 plaque size reflects faster diffusion which could be attributed to its longer adsorption rate allowing wider diffusion before host lysis. In contrast, fBSZT1 has very short adsorption period and high burst size resulting in locally intense but spatially confined lysis.

To determine the phage applicability among APEC, we isolated 51 strains from various Indian states and determined the antimicrobial susceptibility pattern. We observed high resistance to ciprofloxacin (47%), nitrofurantoin (45.1%), colistin (47%), and tetracycline (41.2%) while approximately 30% of strains were resistant to cephalosporins. Notably, 23 strains (45.1%) were resistant to 3 or more antibiotics of different classes, making them MDR. These findings align broadly with global trends but reveal important regional variations. Globally, resistance prevalence of *E. coli* from food animals is 59% against tetracycline, 57% to ampicillin, 35% to chloramphenicol, 30% to ciprofloxacin, 28% to gentamicin, and 33% to cefotaxime (26). Our observed MDR rate of 45.1% is substantially lower than the 94% reported previously in northern India (27), but it still represents a critical public health concern, with nearly half of all isolates resistant to multiple drug classes. This variation underscores the regional diversity of AMR and the need for adaptable solutions like phage therapy.

APEC serotype distribution, varies by geographic region, has important implications for phage therapy development. Globally, O1, O2, and O78 serogroups typically accounts for 80% of APEC isolates, but the prevalence varies with some studies showing these serotypes prevalence to be only 7.7% (1, 28). In our study, we found none of the strains with these serotypes, though other relevant APEC serotypes were observed (29). In a study on APEC in India, O1, O2, and O78 were also not observed, but O26 (32%), O98 (28%), O120 (14%), O11 (12%), O135 (8%), and O17 (4%) serotypes were prevalent. Out of these serotypes, we observed O120 (7.8%), O11 (1.96%), and O17 (3.92%) along with other serotypes O101 (17.65%), O109 (5.88%), O111 (3.92%), O119 (7.84%), O128 (1.96%), O156 (3.92%), O16 (5.88%), O169 (3.92%), O18 (1.96%), O20 (1.96%), O5 (3.92%), O7 (9.8%), O84 (5.88%), and O9 (1.96%). (30). Notably, in Brazil, where the prevalence of O1, O2, and O78 serogroups is low, the MDR prevalence is also lower (38%), which corresponds to our study (1). Whether this correlation reflects an epidemiological linkage between specific serogroups and resistance mechanisms, or regional differences in antibiotic selection pressure, remains unclear. Regardless, these findings emphasize developing effective phage formulations tailored to regional pathogen populations.

Among all APEC isolates, approximately 50% of isolates were sensitive to all phages, while 52.17% of MDR strains were lysed by at least one out of five phages. Our findings are comparable to or exceed previous reports showing the efficacy of phages against APEC both *in vitro* and *in vivo* (1, 15, 31, 32). It is also noteworthy that four of the five phages showed slight activity against *E. coli* laboratory strains such as BL21 and MS145-7 **(Fig. 7)**, indicating that phage receptors are not restricted to APEC and are present on non-pathogenic *E. coli* strains as well. The phages also showed activity against a *Shigella flexneri*, and a *Shigella sonnei* strain. Notably, two of our phages, fUAMT1 and fPKPT2, despite showing >90% intergenomic similarity displayed varying host range. Comparing their tail fiber genes revealed differences in their host-recognition apparatus. The long-tail fiber distal subunit differs by 24 bp, and the accessory host-specificity tail fiber differs by 21 bp among these two phages. Furthermore, the fUAMT1 encodes an additional tail fiber protein which is absent in fPKPT2, potentially expanding its receptor interaction surface. These differences in the tip regions of the long-tail fibers responsible for phage host recognition is a probable molecular basis for the divergent host ranges of fUAMT1 and fPKPT2. The difference in the host range of all 5 phages is likely attributable to a combination of factors like the availability of specific phage receptors on bacterial surface, and post-adsorption intracellular defence systems such as restriction-modification systems, CRISPR-Cas machinery, anti-phage defence islands, and abortive infection mechanisms present in non-susceptible bacterial hosts (33). Distinguishing between these mechanisms was beyond the scope of this study and should be explored in future work.

The use of *G. mellonella* as an infection model provides a valuable bridge between *in vitro* screening and avian trials as it has an innate immune system similar to vertebrates, which maintains its practical and ethical advantages (34). The *in vivo* efficacy of phages in the *G. mellonella* larvae revealed that three phages, particularly fPKPT2 and fUAMT1, can significantly improve the survival of larvae by 80%, confirming the phage’s ability to control systemic infections in an *in vivo* model. Notably, fRPOT1 and fDMYT1 showed reduced *in vivo* efficacy despite achieving comparable *in vitro* activity to effective phages (fBSZT1, fUAMT1, fPKPT2). The differences in the lytic activity of all 5 phages in an *in vivo* model suggest that factors such as phage stability in hemolymph, immune recognition, or spatial distribution may influence therapeutic outcomes. None of the phages achieved 100% larval survival even at MOI 10; efficacy in a more complex avian model may be lower and must be validated before clinical application. This lack in individual *in vivo* phage activity and the emergence of phage-resistant bacteria in the OD-based liquid suppression assay **(Fig. 8)** represents an important barrier to phage therapy. The use of phage cocktails or phage-antibiotic combinations may reduce resistance development and should be prioritised in future studies. The phage and antibiotics showing negative Pearson correlation based on activity could be tested experimentally for their interactions (35).

Several recent studies have shown *in vitro* efficacy of APEC phages, but comprehensive characterization of larger, geographically relevant phage banks remains limited. Recently, a phage cocktail, FÓRMIDA, showing high specificity against APEC and HR APEC pathotypes (1) was developed, but its efficacy in an *in vivo* model is yet to be determined. Notably, the two of our phages showing *in vivo* efficacy, fPKPT2, and fUAMT1, along with phage fRPOT1, can lyse the maximum number of tested APECs individually (64.71%). The combination of these 3 phages is also predicted to be the best combination by the PhageCocktail plugin in Cytoscape (36). While combining phages could kill bacteria more efficiently due to synergism, antagonism can also occur (37). The receptor of each phage could be identified to prepare a phage cocktail with phages targeting different receptors.

In summary, we report a fully characterized new lytic phage bank targeting APEC serotypes from Indian poultry. Our findings support the phages usefulness in future phage cocktail development and suggest a higher dosage for therapeutic application of phages. Encapsulation and freeze drying may enhance the phage stability and delivery to the target site, maintaining the effective phage concentrations over extended periods (35). Further investigation is required to test phage-phage interactions both *in vitro* and *in vivo*, assess resistance development rates, and validate efficacy in chicken infection models before preparing a therapeutic cocktail.

## 4. Materials and Methods

### 4.1. Chemicals, Bacterial Strains, and Culture Conditions

The reagents and culture media were purchased from HiMedia Laboratories (Mumbai, India), and the plastic-wares were obtained from Tarsons (Kolkata, India). The reference bacterial strains were obtained from BEI Resources, NIAID, NIH, USA; Microbial Type Culture Collection and Gene Bank (MTCC), Chandigarh, India; Veterinary Type Culture Collection (VTCC), and National Centre for Veterinary Type Culture (NCVTC), Hisar, India. The APEC strains and bacteriophages were isolated from poultry faecal matter collected from several states of India **(Supplementary Table S1 and S6)**. For all experiments, bacteria were grown in Tryptone Soya Broth (Soybean Casein Digest Medium), Luria Bertani (LB) broth, LB agar, and Soybean Casein Digest Agar (TSA) at 37°C for 18h to 24h.

### 4.2. Bacteriophage isolation and purification

Bacteriophages were isolated from the poultry fecal samples against the host *E. coli* O135 (APEC serotype found in Indian poultry) (30) by the enrichment method (38). For monophage isolation, the filtered supernatant obtained after enrichment was serially diluted and mixed with an overnight-grown bacterial culture, added to the molten soft agar (0.75%), overlaid on the TSA plates, and incubated for 24 h at 37°C for the development of plaques. The single plaque was suspended in SM buffer (50 mM Tris-HCl, 100 mM NaCl, 8 mM MgSO_4_, pH 7.5). To ensure the purity of the bacteriophage, the plaque assay was repeated five times by collecting the consecutively isolated plaques (17, 39).

### 4.3. Propagation and concentration of phages

The 1 mL of an overnight culture of the bacterial host was added to 100 mL of LB medium containing 10 mM MgSO_4_ and 10 mM CaCl_2,_ and incubated at 37°C with vigorous shaking (190 rpm) till the culture reached the exponential phase (OD_600_ = 0.7). Then the bacteriophage lysate at a MOI between 0.1–1 was added and incubated for 6 h, shaking vigorously (190 rpm) at 37°C until the bacterial growth inhibition observed in comparison to uninfected control. The lysate was then separated into 50-mL centrifuge tubes and centrifuged at 60000g for 10 min at room temperature. The supernatant was then filtered out with a 0.22 µm filter and stored at 4°C. After that, to further remove the debris and endotoxins, the phage lysate was treated with chloroform and octanol, respectively (40). The supernatant was concentrated by treating it with 20% polyethylene glycol (PEG 8000) (Sigma, USA) in 2.5 M NaCl. The pellet was dissolved in SM buffer and then stored for further experiments (40, 41).

### 4.4. Determination of phage titer

The titer of the lysate was determined through the double-layer agar overlay (DLA) method (42). The bacteriophage lysate was serially diluted (10^-1^ to 10^-8^) in SM buffer to obtain a dilution with countable plaques. Then the diluted bacteriophage lysate was added to 3 mL of top agar and 500 µL of host bacterial culture mixture and overlaid on TSA plates. The plates were incubated at 37°C overnight, and bacteriophage plaques were counted. To calculate the titer in plaque-forming units (PFU/mL), the following formula was used:

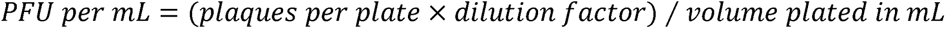

### 4.5. Plaque size and bacteriophage morphology through transmission electron microscopy (TEM)

The plaque morphology of the individual bacteriophages was tested with the host *E. coli* strain. The DLA overlay method was used to determine the plaque size, morphology, and diameter (42, 43). The TEM was performed to examine the structure of bacteriophages. The purified bacteriophage lysate was centrifuged at 20,0000g for 90 min and washed with 0.1 M ammonium acetate (pH 7.0). A drop of bacteriophage lysate was placed on the copper grid, negatively stained with 2% (m/v) uranyl acetate for 30s, dried with filter paper, and then observed under Tecnai G20 HR-TEM at 200kV at Sophisticated Analytical Instrumental Facility (SAIF), AIIMS, New Delhi.

Based on the TEM morphology, the bacteriophage class was identified and classified (44, 45). The size of phages was measured using ImageJ 1.54p (NIH, USA) (46), and the mean and standard deviation of head and tail length were calculated for at least 10 phages.

### 4.6. Stability of phages at different temperatures and pH conditions, and long-term storage

While the phage treatment given to chickens requires phage stability at the physiological level, its storage is maintained at lower temperatures. To determine bacteriophage stability when exposed to low or high heat and pH, bacteriophages were exposed for one hour to a range of temperatures of 4, 15, 25, 37, 45, 50, 55, 60, 65, 70 and 80°C or diluted in SM buffer (1:10) of pH 2, 3, 4, 5, 6, 7, 8, 9, 10, 11, and 12. To determine bacteriophage titer after exposure to different temperatures and pH conditions, the bacteriophage lysates were serially diluted, and the DLA overlay method was performed (39). The phages were kept at 4°C, -20°C, and -80°C for storage. The phages kept at -20°C and -80°C were stored in 15% glycerol (prepared in SM buffer). The phage titers were calculated for stored phages after every 6 months to determine their long-term stability under various storage conditions (47).

### 4.7. Phage adsorption and one-step growth curve assay

Adsorption efficiencies of bacteriophages were determined in replicates as described previously (48). In brief, suspensions of bacteria and bacteriophages at MOI 1 were incubated in 5 mL double strength (2Χ) TSB. The volume of the solution was made up to 10 mL by adding SM buffer and then incubated at 37°C for 25 min. After every 2 min up to 10 min and then after every 5 min, 100µL suspension was collected and centrifuged at 80000g for 1 min to pellet bacteria and adsorbed bacteriophages. The supernatant was used to perform the DLA overlay to determine the PFU/ml of un-adsorbed phages.

The one-step growth experiment was performed by taking the host *E. coli* culture with 10^7^ CFU/mL and mixing it with bacteriophage at an MOI of 0.01 (49). Bacteriophages were adsorbed for 15 min at 37°C, and unbound bacteriophages were removed by centrifuging at 80000g for 5 min. The bacterial pellet was resuspended in TSB and incubated at 37°C. Aliquots were taken every 10 minutes for 1.5 h, and the DLA overlay was performed to determine the PFU/mL count. The latent period was read directly, and the burst size of bacteriophage, which is the ratio of the final count of liberated bacteriophage particles to the initial amount of infective bacterial cells, was calculated as described previously (50, 51).

### 4.8. Identification of putative phage receptor types on the bacterial surface

To determine whether the phage is binding to the outer membrane protein (OMP) or carbohydrate, the host *E. coli* was treated with either proteinase K or sodium periodate as described earlier (52). Briefly, 1 mL of overnight-grown bacterial culture was pelleted, and either dissolved in 0.5 mL TSB and treated with 5 µL proteinase K (20 mg/mL; HiMedia) for 3 h at 37°C or suspended in 1 mL of 100 mM sodium periodate (Sigma-Aldrich, MO, USA) and incubated for 2 h in the dark at room temperature. The sodium periodate solution was prepared in sodium acetate; therefore, only sodium acetate was added in the control. Following incubation, the bacterial culture was pelleted, washed with TSB, resuspended in 900 µL TSB, and treated with the respective phage at MOI 1 for 1 h at 37°C. Then the number of non-adsorbed phages was measured after centrifugation using the DLA method.

### 4.9. APEC isolation and identification

The chicken droppings, both from infected broilers and layers, were collected from commercial poultry farms and small holder farms across India in sterile containers individually **(Supplementary Table S6)** and transported to the laboratory. We collected samples from 5 states: Odisha (n=8), Bihar (n=7), Uttar Pradesh (n=10), Punjab (n=6), Rajasthan (n=18), and 1 union territory: Delhi (n=2), and processed within 48 h of collection. 1 g of each sample was seeded in 10 mL LB broth and incubated for 30 min at 37°C and 140 rpm for the release of bacteria. The *E. coli* strains were isolated by culturing sample dilutions on MacConkey agar plates. Based on morphological appearance and on comparison with a positive control, *E. coli* DH5α (MTCC 1652), suspected *E. coli* colonies were selected. The *E. coli* strains were then confirmed with Gram staining, biochemical assays such as oxidase, urease, Triple sugar iron (TSI), and citrate tests, and outsourced for serotyping through the **<u>C</u>**entral **<u>R</u>**esearch **I**nstitute (CRI), Kasauli, India. Finally, the confirmed isolated bacteria were stored in 25% glycerol at -80°C. Further, a PCR with primers *hly*F, *omp*T, *iro*N, *iut*A and *iss* was performed to confirm APEC phenotype (21, 53). The PCR products were observed on a 1.5% agarose gel stained with SYBR Safe DNA Gel Stain (Invitrogen, USA) **(Supplementary Table S8).**

### 4.10. Antibiotic susceptibility testing (AST) of APEC isolates

The AST of the APEC isolates was performed using the Kirby-Bauer disk diffusion assay. The isolates were screened against 10 commercially available antibiotics from various classes and different generations, which are used for veterinary and human medicine purposes, to find the resistance prevalence and pattern in the poultry sector in India. Antibiotics such as cephalosporins (ceftazidime-CAZ (30µg), cefuroxime-CXM (30µg)), carbapenems (Ertapenem-ETP (10µg)), penicillins with beta-lactamase inhibitor (amoxicillin/clavulanic acid-AMC (10µg), piperacillin/tazobactum-PIT (100/10µg)), nitrofurans (nitrofurantoin-NIT (300µg)), polymyxins (colistin-CL (10µg)), fluoroquinolones (ciprofloxacin-CIP (5µg)), tetracyclins (tetracycline-TE (30µg)), and sulfonamides (cotrimoxazole-COT (25µg)) (54) were used to observe susceptibility. All the antibiotic disks used were purchased from HiMedia, India. The AST was performed on Muller-Hinton agar (MHA) plates (HiMedia) with the concentration of freshly grown bacteria equal to 0.5 McFarland units as per the Clinical and Laboratory Standards Institute (CLSI) guidelines (55). Further, susceptibility of 22 strains having higher resistance was checked with 8 more antibiotics belonging to nitrofurans (nitrofurazone-NR (100µg), and furazolidone-FR (100µg)) and fluoroquinolones (gemifloxacin-GEM (5µg), pefloxacin-PF (5µg), ofloxacin-OF (5µg), levofloxacin-LE (5µg), norfloxacin-NX (10µg), and moxifloxacin-MO (5µg)).

### 4.11. Bacteriophage DNA isolation

To remove residual bacterial DNA and RNA present in the concentrated bacteriophage lysate, 425 µL of concentrated filter-sterile phage lysate was incubated with 5 µL DNase-I (stock at 1 mg/mL), 50 µL of 10X DNA digest Buffer, and 20 µL of RNase-A enzyme (stock at 10 mg/mL) for two hours at 37°C without shaking. The lysate was cooled down at room temperature and stored at 4°C (40, 41). Phage genomic DNA from the lysate was isolated using the PCI method (56). The 1.25 µL of Proteinase K (20 mg/mL), 0.5% SDS, and 20 mM EDTA (pH 8.0) were added to the DNase-treated lysate and incubated for 1 h at 60°C. Then, a mixture of **<u>p</u>**henol, **<u>c</u>**hloroform, and **i**soamyl alcohol (PCI) (25:24:1 v/v/v) (HiMedia) was added to the sample and mixed thoroughly, centrifuged for 10 min at 13,000×g, and the aqueous phase was collected in a sterile tube. 1/10 volume of 3 M NaOAc (pH 5.2) and 2.5 volumes of ice-cold ethanol were added to the supernatant and incubated at -20°C overnight to precipitate the DNA. Then it was centrifuged for 20 min at 15,000×g, and the supernatant was discarded entirely. After that, the pellet was washed with 70% ethanol twice, dried, and resuspended in 20μl of nuclease-free water. The DNA quantification was performed using a Tecan Spark® multimode microplate reader, and the quality check of DNA integrity was done by gel electrophoresis on a 0.8% agarose gel.

### 4.12. Whole genome sequencing, assembly, and annotation

Whole-genome sequencing of the bacteriophage was performed at Unipath Laboratories, Ahmedabad, India. The DNA quantity was checked with Qubit® 4.0 fluorometer, and a paired-end sequencing library was prepared using the Twist NGS library preparation kit for Illumina® (Cat no/ID 104119). The amplified libraries were analyzed on Agilent TapeStation 4150 using high-sensitivity D1000 ScreenTape® as per the manufacturer’s instructions. The libraries were then loaded onto the Illumina Novoseq X Plus platform for cluster generation and sequencing using 2x150 bp chemistry (17). The quality of the raw sequence data was determined with FastQC v0.12.1 (57). The low-quality genomic reads were removed with Trim Galore 0.6.4 (58), and the genomes were assembled using MetaViral SPAdes 3.15.4 (59). The assembly elements were determined with Unigenome’s Perl script, and the best scaffold for each phage was considered for downstream analysis. The final genome assemblies were run through checkV 1.0.3 (60), tRNAscan-SE 2.0 (61), and Prodigal 2.6.3 (62) for determining completeness, predicting tRNA and genes, respectively. The predicted genes were run with BLASTP 2.13.0+ (63) for similarity search against NCBI’s non-redundant (NR) and CARD (64) databases to identify function and AMR genes, respectively. To increase confidence in the predicted ORFs and CDSs, data were compared to those expected through PHASTEST 1.0.1 (65) and Prokka 1.2.0 server (accessed through Galaxy version 1.0.1, https://usegalaxy.org/) (66, 67). Further, Blast2GO 1.5.1 was used for gene ontology (GO) mapping and annotation (68). The raw data and final assembled genomes were submitted to GenBank **(Supplementary Table S1)**. For mass screening of genes for antimicrobial resistance (AMR) or virulence genes, the gene nucleotide sequences were run through ABRicate 1.0.1 and searched against CARD, ecoh, ecoli_vf, megares, ncbi, plasmidfinder, resfinder, and VFDB databases with a minimum 80% DNA coverage and 80% DNA identity (69).

### 4.13. Bioinformatics analysis of phage genomes

A genomic map for each phage was constructed using the Proksee CGview program Map Builder 2.0.5 (https://proksee.ca/). GC content and skew were added in the map using GC content 1.0.2 and GC Skew 1.0.2 tools, respectively (70). Annotations were incorporated in the map using Phigaro 1.1.0 (71), mobileOG-db 1.1.3 (72), PHASTEST 1.0.1 (65), Prokka 1.2.0 (67), and VirSorter 1.1.1 (73). The lifestyle of each phage was identified with Life Cycle Classifier v1.6.0 accessed through PhageAI 1.0.2 (https://www.phage.ai/) (18). A proteomic tree was generated in ViPTree version 4.0 (https://www.genome.jp/viptree/) based on genome-wide sequence similarities computed by tBLASTx with reference viral genomes (74). Further, the similarity scores were identified, and each phage was aligned with its closely related phage by DiGALign version 2.0 (75).

VIRiDIC applies BAVS traditional algorithms to calculate phage genome similarities (19). The pair-wise inter-genomic similarities between phages were evaluated with the Virus Inter-genomic Distance Calculator (VIRIDIC) (http://rhea.icbm.uni-oldenburg.de/VIRIDIC/) as recommended by ICTV (International Committee on Taxonomy of Viruses), subcommittee of Bacterial and Archaeal Viruses (BAVS). Phage taxonomy according to ICTV was identified with taxMyPhage 3.3.6 (https://ptax.ku.dk/) (76).

A genome-based phylogeny and classification were performed with the VICTOR tool (https://victor.dsmz.de) (20). The comparisons of nucleotide sequences were conducted using the Genome-BLAST Distance Phylogeny (GBDP) method (77) under settings recommended for prokaryotic viruses (20). The resulting inter-genomic distances were used to infer a balanced minimum evolution tree with branch support via FASTME, including SPR post-processing (78) for formula D0. Branch support was inferred from 100 pseudo-bootstrap replicates each. Trees were rooted at the midpoint and visualized with ggtree (79). Taxon boundaries at species, genus, and family level were estimated with the OPTSIL program (https://cran.r-project.org/web/packages/optpart/optpart.pdf) using the recommended clustering thresholds and an F value (fraction of links required for cluster fusion) of 0.5.

To further strengthen the phylogenetic relation and classification of phages, two core genes, major capsid protein (MCP) and terminase large subunit (TerL), were used to generate a neighbour-joining tree. The amino acid sequences of these two proteins were aligned to already available bacteriophage proteins identified as the closest type species by BLASTp against the NCBI nucleotide database, having over 90 percent identity. The muscle alignment was performed using MEGA version 11.0.13 (80), then the phylogenetic analyses were conducted, and the evolutionary history was inferred using the Neighbor-Joining method. The bootstrap consensus tree inferred from 1000 replicates was taken to represent the evolutionary history of the taxa analyzed, showing evolutionary distances computed with the Poisson correction method.

### 4.14. Host range determination and specificity

The bacteriophage activity was assayed through spot assay against the isolated MDR APEC isolates. The zone of clearance observed in the 6 h spot test was used to determine host sensitivity. It was scored as complete lysis (+++), moderate lysis (++), turbid lysis (+), and no lysis (-) (39). The phage lytic ability against all bacteria was analyzed with the help of Cytoscape 3.10.3 plugin app PhageCocktail 1.1 (36). The activity of the isolated bacteriophage against different laboratory strains was also determined by spot assay method described above. Various reference bacteria such as *Escherichia coli* MS145-7 (ATCC, HM-341), E. *coli* BL21 (DE3) (MTCC 1679), *E. coli* O:108 (VTCC BAA462), *E. coli* DH5α (MTCC 1652), *Shigella* D9 (ATCC, HM-87), S. *flexneri* 24570 (ATCC, NR-517), S. *boydii* NCTC 12985 (ATCC, NR-521), *S. sonnei* WRAIR (ATCC, NR-519), *Salmonella enterica subspecies enterica* S5564 (serovar Typhimurium) (ATCC, NR-22070), *Enterococcus faecalis* Strain TX2137 (ATCC, HM-432), *Staphylococcus aureus* Strain HIP13170 AKA VRS3a (ATCC, NR-46412), *Mycobacteria smegmatis* (MTCC 992), and *Bacillus subtilis* 168 (ATCC, NR-607) were plated on the TSA plates and 10 µL spots of bacteriophages (10^8^ PFU/mL) were added to determine the host range of new phages. Based on the bacteriophage activity against the APEC isolates and the AST pattern of each isolate, the Pearson correlation among the bacteriophages and antibiotics was determined. The correlation coefficient r was determined with the help of GraphPad Prism.

### 4.15. Lytic efficiency of *Escherichia* phages at different MOIs

A liquid suppression assay was performed to determine the effect of phages against *E. coli* for 48 h (81). Briefly, an overnight culture of *E. coli* O:135 was diluted to 2×10^6^ CFU/mL in 2Χ TSB and added to a 96-well microtiter plate. An equal volume of each phage diluted in SM buffer was added in triplicate wells at MOIs of 0.1, 1.0, and 10.0. The plates were incubated at 37°C with continuous shaking at 150 rpm, and absorbance (OD_600_) was measured at different time intervals up to 48 h.

### 4.16. In vivo efficacy in Galleria mellonella

The *G. mellonella* larvae were used to determine the effect of phage in preventing *E. coli* infection, in accordance to current institutional guidelines. Prior to performing experiment, larvae were starved for 24 h. Initially, to determine the effective dose of bacteria, the healthy larvae weighing 250 ± 25 mg were washed with 70% ethanol and separated into groups of 10 per set. After ethanol washing, larvae were kept at room temperature for 2 h, the dead larvae were removed and only live were used for experimentation. In each set, 10 µL of an overnight-grown host *E. coli* O135 was injected into the last left proleg of the larvae, with the final bacterial concentrations ranging from 10^2^ CFU/mL to 10^6^ CFU/mL diluted in PBS. Only PBS was used as a negative control in one set. The larvae were placed in the dark at 37°C for 5 days without food, and mortality was observed for 5 days. The larval movement and melanization were considered parameters for their mortality. The melanised larvae without movement were recorded as dead. For larvae rescue by phage, a rescue experiment was performed where 10 µL of bacteria was injected into the last left proleg at the final concentration of 5Χ10^5^ CFU/mL. At 1 h post-infection, 10 µL of phage in PBS at MOI 10 (5Χ10^6^ PFU/mL) was injected into the last right proleg. 5 different sets were used for 5 phages, and a set of controls was injected with PBS alone. In each set 5 larvae were injected. In negative controls, the bacteria were replaced by PBS, and in one set of phage-only controls, a phage cocktail was injected. The larvae’s survival and melanization were recorded for 5 days. At the end of the experiment, the larvae were crushed, homogenized in PBS, serially diluted, and plated on MacConkey agar plates to determine CFU/mL (39, 82).

### 4.17. Statistical Analysis

All experiments were performed in triplicate, and the quantitative data obtained were analyzed. A t-test was performed using GraphPad Prism, version 8 (https://www.graphpad.com/scientific-software/prism/). A log-rank test was performed to compare the survivability of *G. mellonella* larvae within the groups studied. The comparisons were made with respective controls, and the data were plotted with mean and standard deviation. The significance was recorded as \**p<*0.05, \*\**p<*0.01, \*\*\**p<*0.001, and \*\*\*\**p<*0.0001.

## 5. Conclusion

Our study presents fully characterized new lytic *Escherichia* phages with therapeutic potential against MDR-APEC strains prevalent in Indian poultry. Our five phages possess favourable safety profiles, physicochemical stability, and proven *in vivo* efficacy, with lytic activity against nearly two-thirds of tested isolates. While challenges like resistance evolution, determination of exact receptors, cocktail development and avian model validation remain, these phages represent a valuable resource for developing region-specific therapeutic interventions. These characterized phages provide a foundation for developing treatment for colibacillosis in poultry, and it is a step forward in preventing AMR, which is a problem for public health and the environment. Implementing phage therapy in poultry farming could limit antibiotics dependency and prevent environmental pollution, therefore protecting both humans and the ecosystem, these all being goals of One Health framework.

## 6. Data availability

The complete genome sequences of all five bacteriophages characterized in this study have been deposited in the NCBI GenBank database under the accession numbers PQ724394, PV467748, PV467749, PQ724392, and PV419677. Raw sequencing data have been submitted to the NCBI Sequence Read Archive (SRA) under BioProject ID PRJNA1098433. All other relevant data supporting the findings of this study are available from the corresponding author upon reasonable request.

## 7. Research approval

The study was approved by the Central University of Punjab institutional biosafety committee with IBSC certificate number (CUPB/IBSC/2022/13).

## Author contributions

Tushar Midha and Vishakha performed experiments. TM analyzed the results and wrote the first draft of the manuscript. Somesh Baranwal conceived, provided resources, designed, and supervised the study. All authors reviewed, edited, and approved the final manuscript.

## Acknowledgment

We acknowledge the support provided by staff at Unigenome, Unipath Specialty Laboratory Ltd., Ahmedabad, for whole-genome Illumina sequencing, Central Research Institute (CRI), Kasauli, for serotyping, and NCVTC, Hisar, for providing the host bacterial strain, *E. coli* O135. We thank Vyshnav PV, Rashmi Deora, Satyarth Pandey, Shreevar, and Subodh Pradhan for their help with the sample collection. The authors also acknowledge the technical support provided by the Sophisticated Analytical Instrumentation Facility (SAIF), AIIMS, Delhi, for transmission electron microscopy (TEM); National Bureau of Agricultural Insects Resources (NBAIR) for providing *Galleria mellonella* larvae, and BEI Resources, NIAID, NIH for providing reference strains.

