## Supplementary figures for "Genomic characterization and therapeutic potential of five broad-spectrum lytic bacteriophages against multidrug-resistant avian pathogenic *Escherichia coli* (APEC)"

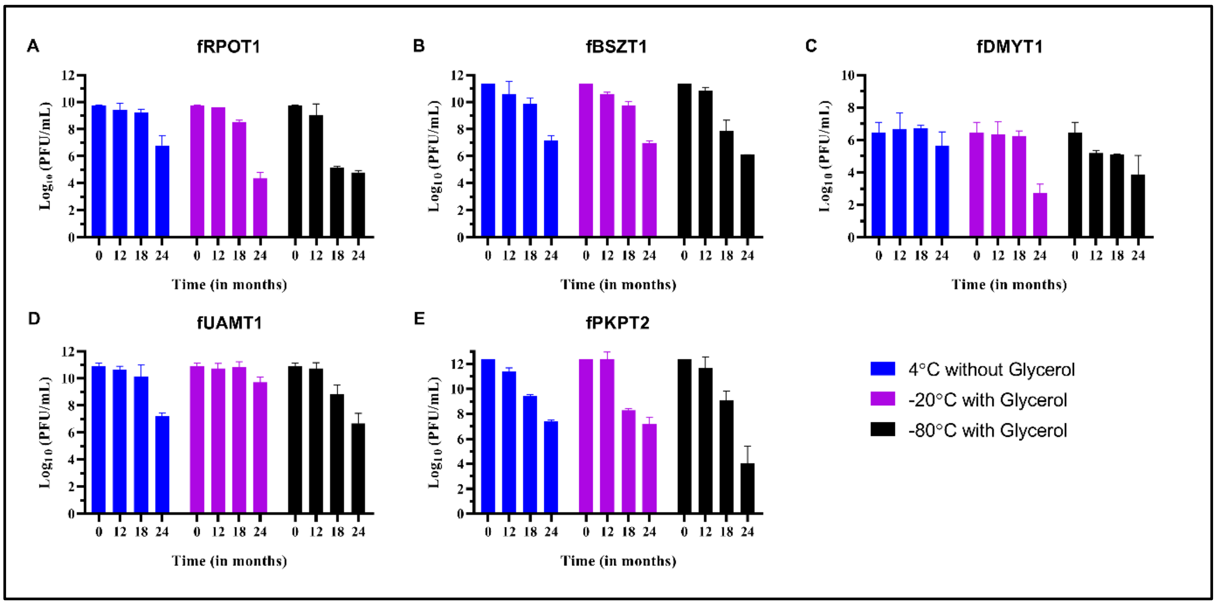


**Fig. S1: Long-term storage of *Escherichia* phages.** The *Escherichia* phage **(A)** fRPOT1, **(B)** fBSZT1, **(C)** fDMYT1, **(D)** fUAMT1, and **(E)** fPKPT2 were stored at 4°C (blue), -20°C (purple) and -80°C (black), and their PFU/ml was determined every 6 months. The values are plotted in $log_{10}$ (PFU/ml) depicting the log reduction phage titer.

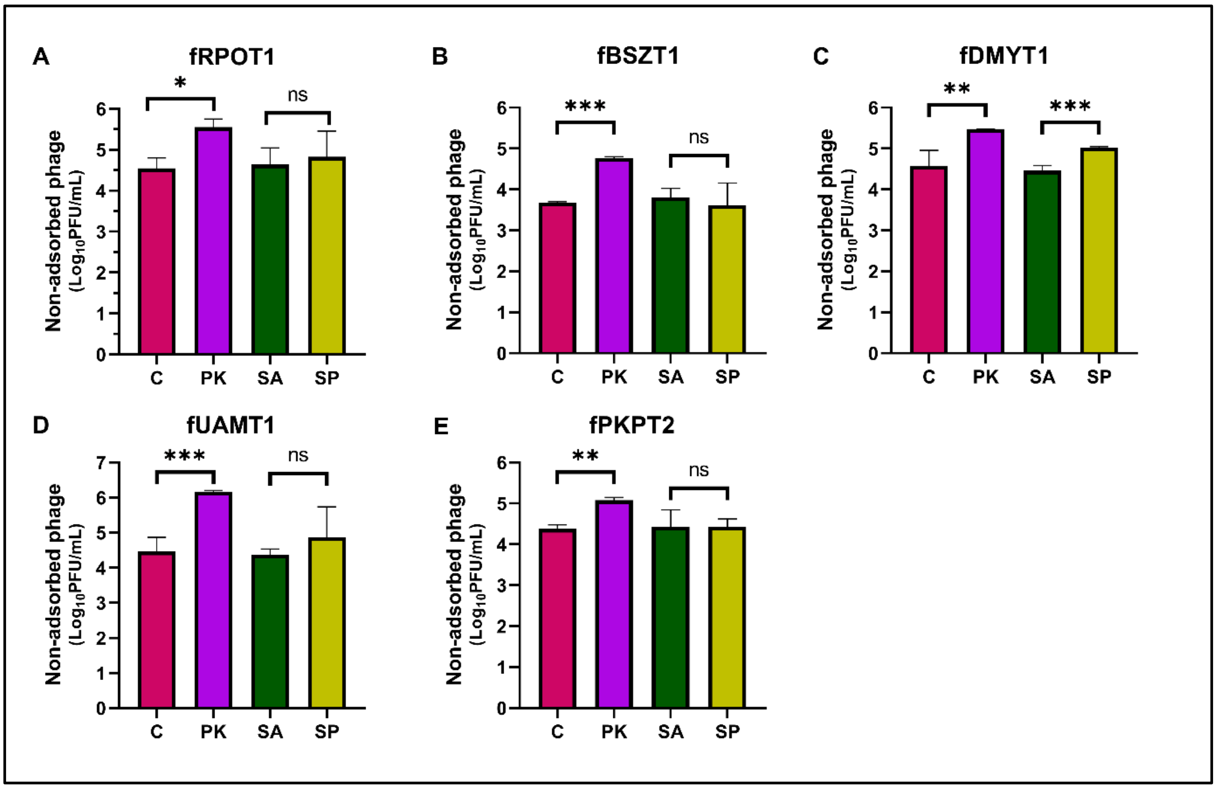


**Fig. S2: Identified receptor type for *Escherichia* phages.** The lipopolysaccharides (LPS) and outer membrane proteins (OMPs) of host *E. coli* were inactivated with sodium periodate (SP) and proteinase K (PK), respectively. For the former, sodium acetate (SA) was taken as a control. The phage adsorption on the treated cell is depicted for phage **(A)** fRPOT1, **(B)** fBSZT1, **(C)** fDMYT1, **(D)** fUAMT1, and **(E)** fPKPT2. For all 5 phages, a significant decrease in phage adsorption is observed upon PK treatment, while for phage fDMYT1, the phage adsorption decreases upon SP treatment as well (D). A t-test is performed to compare the non-adsorbed phages with significance reported as *$p$<0.05, **$p$<0.01, and ***$p$<0.001.

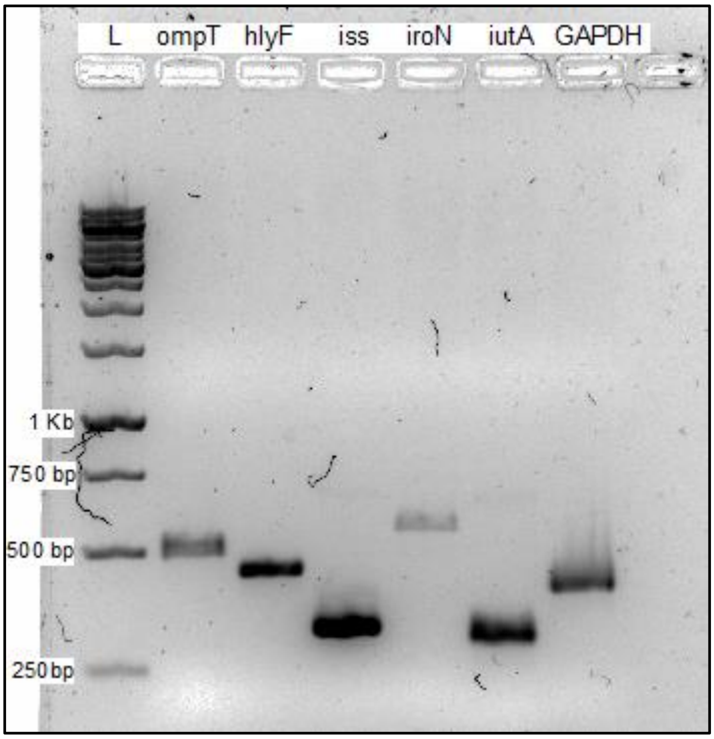


**Fig. S3: Representative image of PCR performed with APEC virulence genes.** PCR was performed with 5 virulence genes of APEC; an episomal outer membrane protease gene, *omp*T; a putative avian hemolysin, *hly*F; episomal increased serum survival gene, *iss*; Salmochelin siderophore receptor gene, *iro*N; and an aerobactin siderophore receptor gene, *iut*A. A bacterial GAPDH gene was used as a positive control.

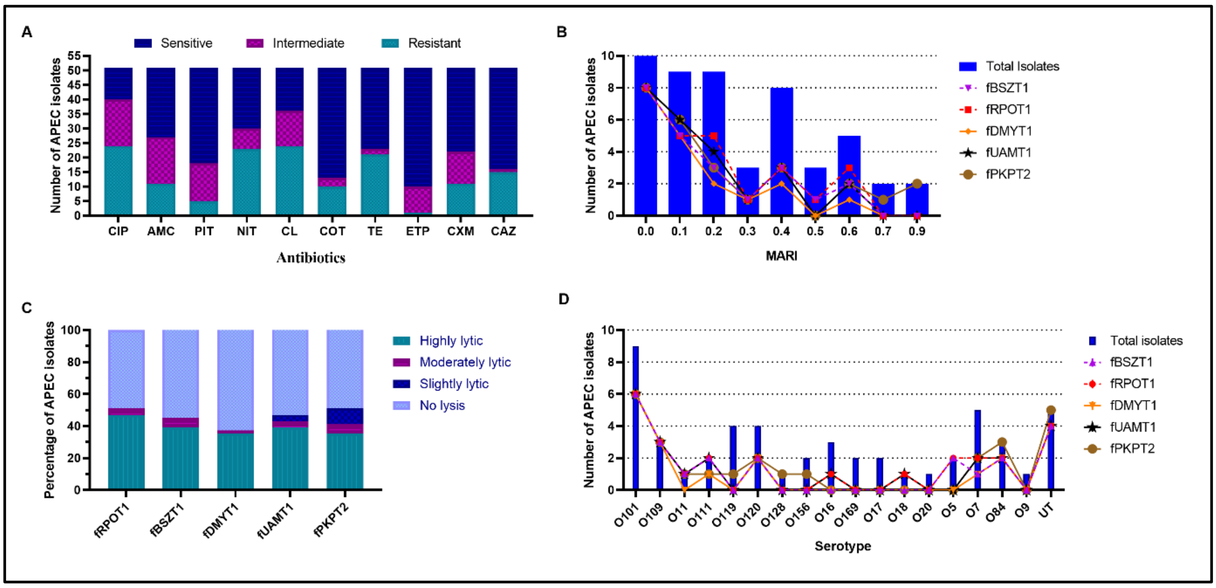


**Fig. S4: Phage activity against MDR-APECs. (A)** Antimicrobial susceptibility testing (AST) of 51 APEC against 10 different antibiotics. **(B)** Based on AST, MARI is calculated for 51 isolates. The number of bacterial strains with a particular MARI is depicted in a bar graph, while the activity of each phage against these strains is shown with a line plot. **. (C)** Percentage of APECs susceptible to 5 *Escherichia* phages is depicted with the level of phage activity shown as per legend (n=51). **(D)** The number of isolates with a specific serotype identified and the activity of each phage against strains of a particular serotype.

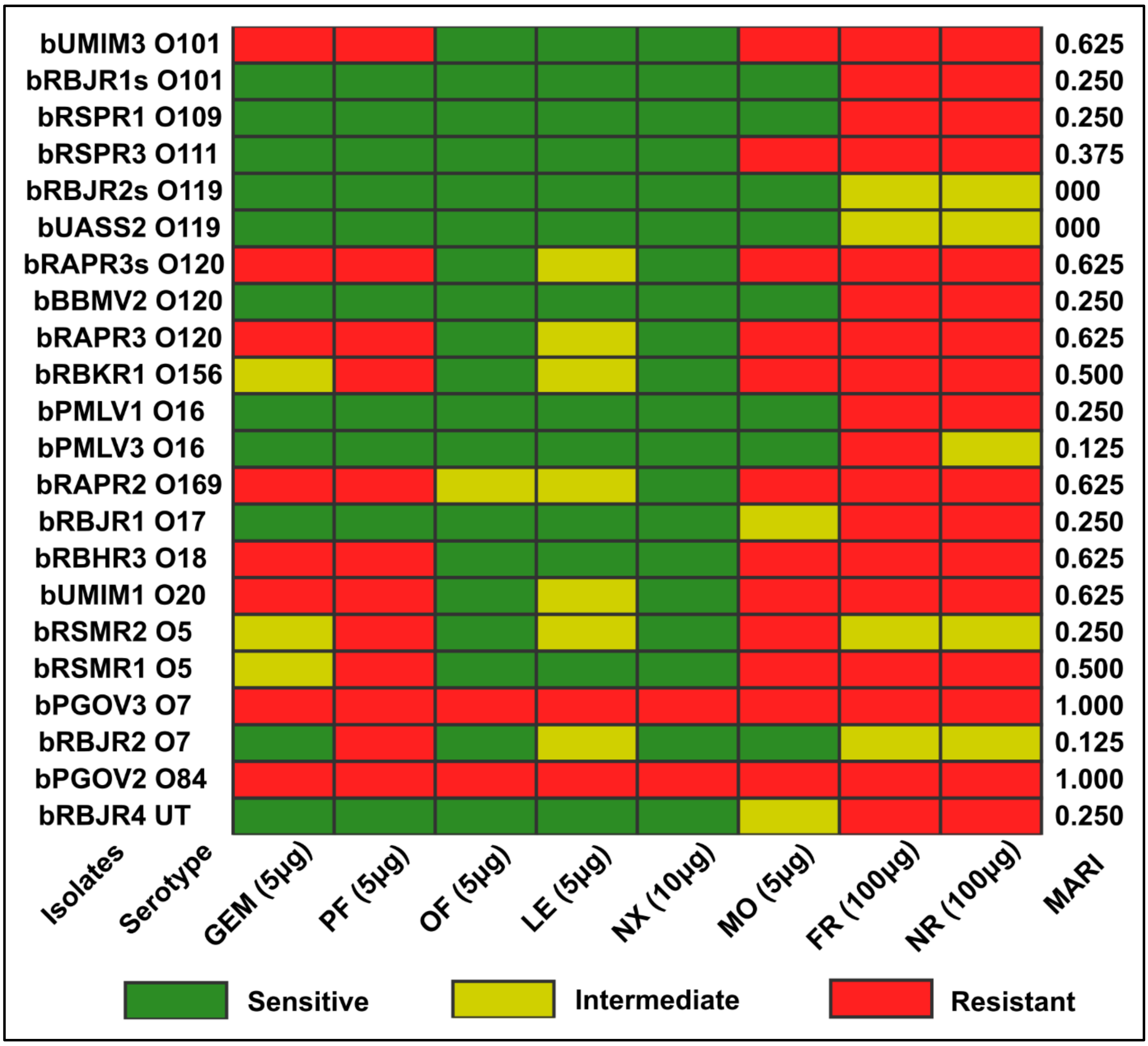


**Fig. S5: Antibiotic susceptibility test of selected 22 strains against 8 more antibiotics.** The bacteria's sensitivity or resistance is depicted in different colors as per the legend. The antibiotics used are gemifloxacin (GEM), pefloxacin (PF), ofloxacin (OF), levofloxacin (LE), norfloxacin (NX), moxifloxacin (MO), nitrofurazone (NR), and furazolidone (FR). The concentration of antibiotic discs used is mentioned on the x-axis along with the respective antibiotics.

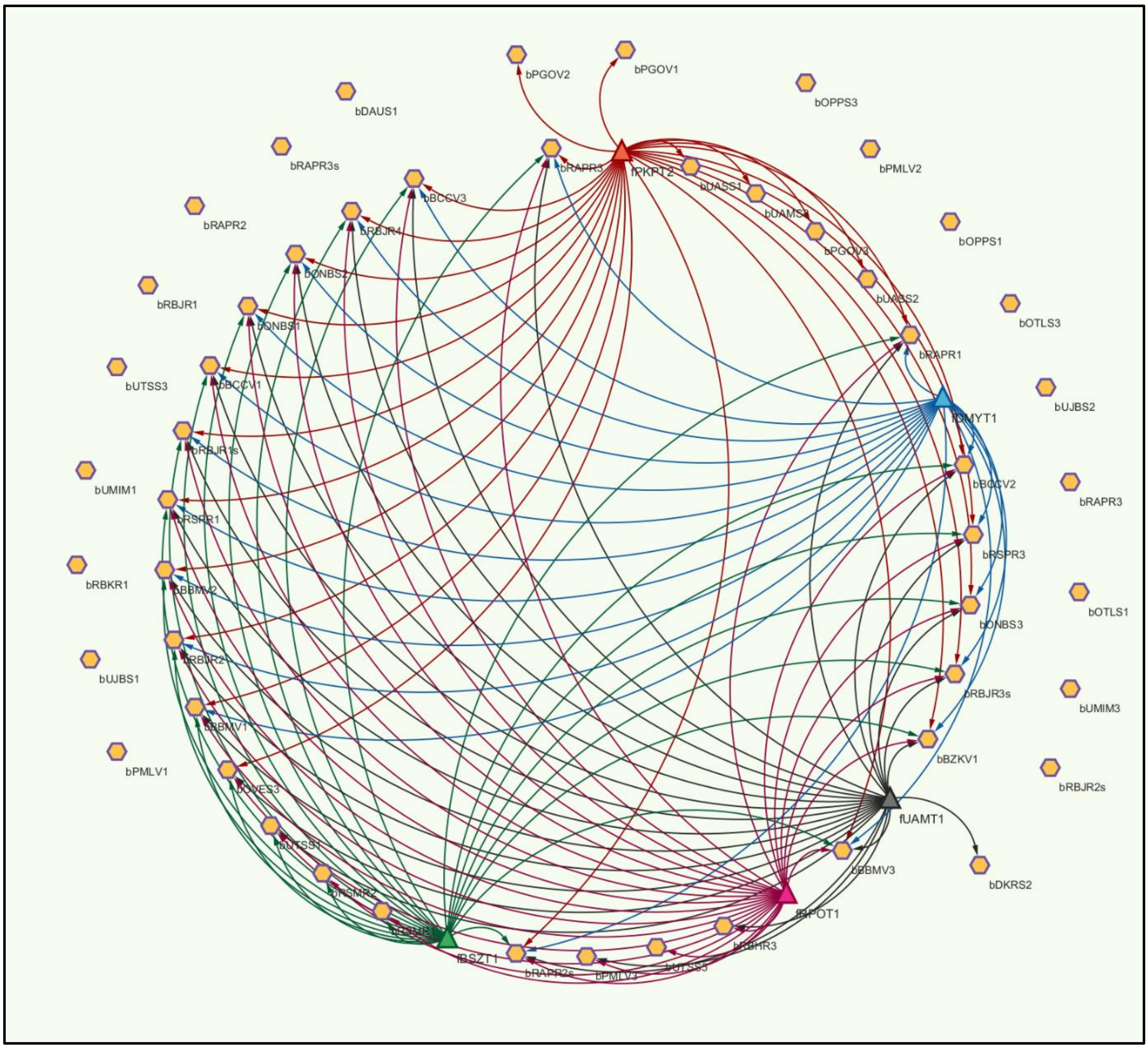


**Fig. S6: Interaction between the 5 bacteriophages and 51 APEC isolates.** The 5 phages and their lytic activity against 51 APEC isolates is depicted with the help of Cytoscape 3.10.3. The bacteria are depicted in a hexagon, and phages in a triangular shape, with different colors showing the interaction of different phages with all bacteria. Out of 51 isolates, 18 isolates were not lysed by any of the phage (outside nodes). Phage fPKPT2 can lyse the maximum number of bacteria (n=26). As many as 33 isolates can be lysed by a combination of 3 phages: fPKPT2, fUAMT1, and fRPOT1.

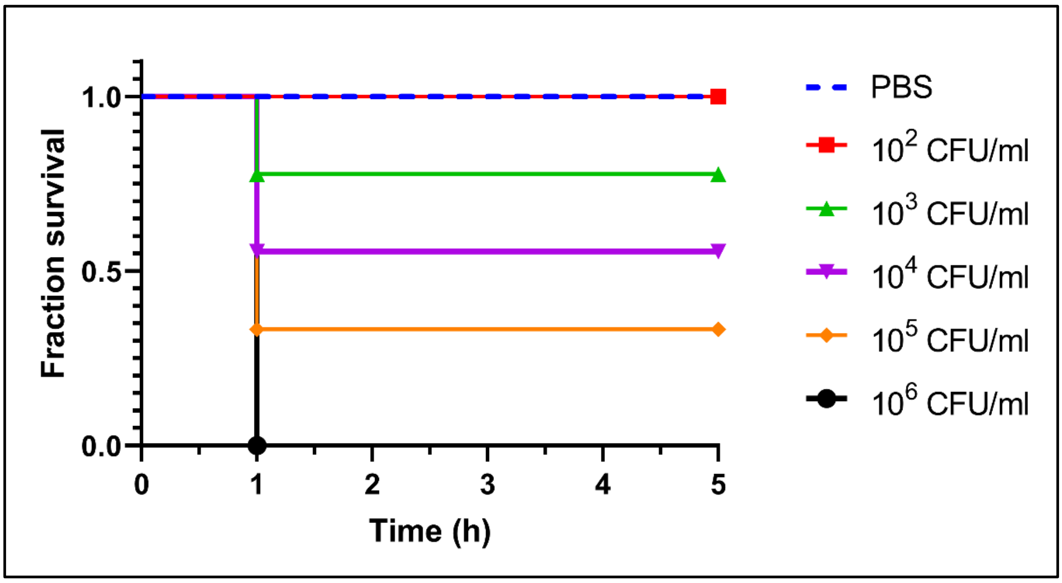


**Fig. S7: Dose-dependent survival curve of *G. mellonella* larvae upon infection with *Escherichia coli*.** The larvae were injected with *E. coli* at different concentrations of $10^2$, $10^3$, $10^4$, $10^5$, and $10^6$ CFU/ml and observed for 5 days. The melanized larvae without any movement were considered dead and recorded.

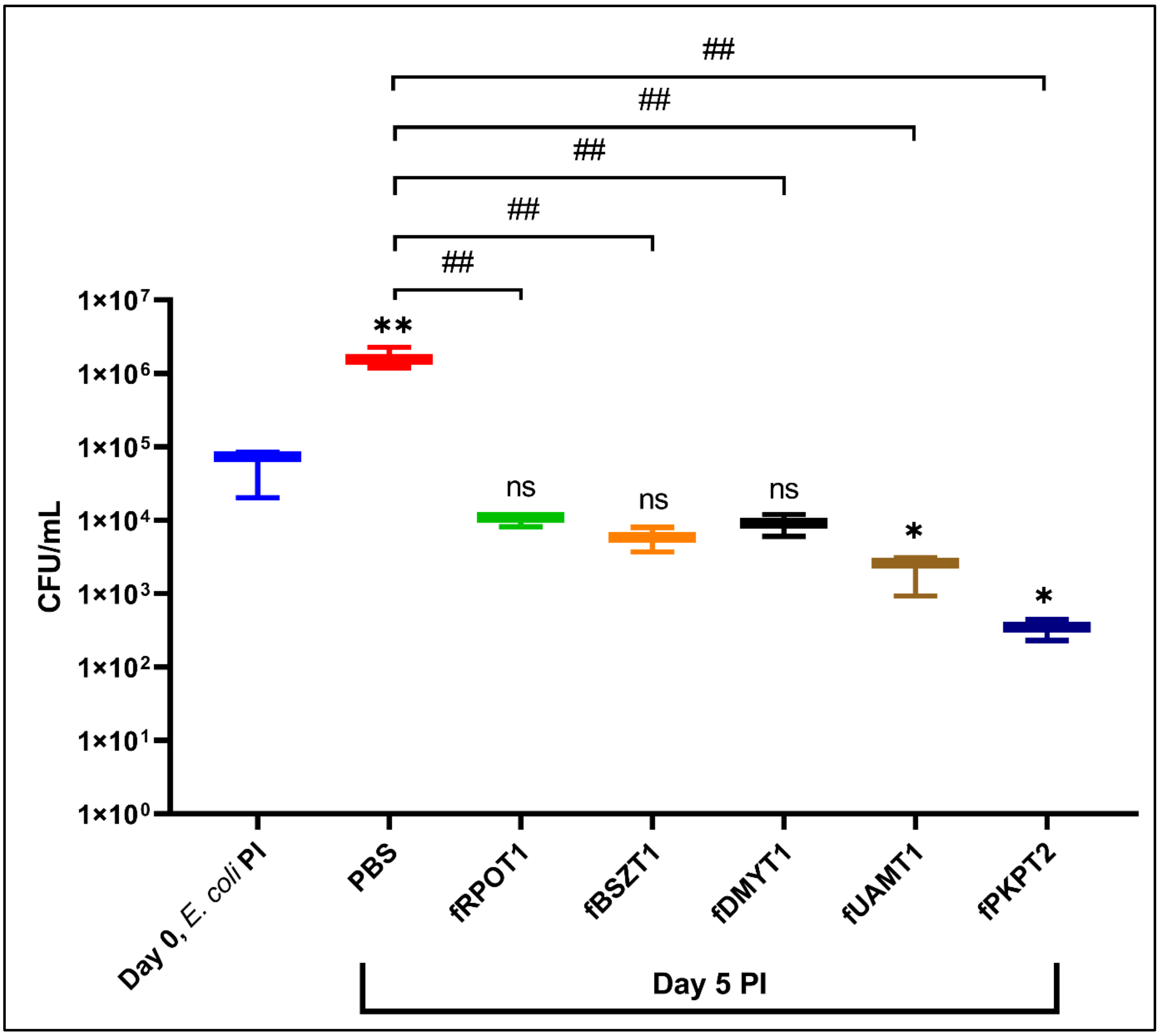


**Fig. S8: Concentration of *E. coli* cells (CFU/ml) inside *G. mellonella* larvae at the time of infection (Day 0) and 5 days postinfection (PI).** In the control set, PBS was injected post 1h infection with *E. coli,* while in other sets, phages were injected at an MOI of 10. A t-test was performed to compare the CFU/ml of each set with the control. The significance of the difference in CFU/ml calculated with the initial injected CFU/ml is depicted in *****, while in comparison to the day 5 CFU/ml of the control is depicted in **#.** (*$p<0.05$; **$p<0.01$, ##$p<0.01$)
