## Supplementary tables for "Genomic characterization and therapeutic potential of five broad-spectrum lytic bacteriophages against multidrug-resistant avian pathogenic *Escherichia coli* (APEC)"

**Table S1:** Properties of five *Escherichia* phages along with the details of samples from which they are isolated.

| Phage | *Escherichia* phage fRPOT1 | *Escherichia* phage fBSZT1 | *Escherichia* phage fDMYT1 | *Escherichia* phage fUAMT1 | *Escherichia* phage fPKPT2 |
|---|---|---|---|---|---|
| ***Sample details*** | | | | | |
| **State/UT** | Rajasthan | Bihar | Delhi | UP | Punjab |
| **Date of collection** | 02-10-2022 | 21-07-2022 | 15-07-2022 | 21-09-2022 | 13-08-2021 |
| **Latitude** | N 25° 4' 10.5" | N 26° 38' 37.9" | N 28° 43' 22.7" | N 25° 30' 46.5' | N 30° 05' 8.5" |
| **Longitude** | E 72° 57' 47.9" | E 84° 54' 14.5" | E 77° 16' 23.6" | E 82° 13' 06.3" | E 74° 50' 31.3" |
| ***Morphological characteristics*** | | | | | |
| **Plaque Size (mm)** | 0.37 ± 0.03 | 0.22 ± 0.03 | 2.14 ± 0.13 | 0.88 ± 0.1 | 0.87 ± 0.08 |
| **Head length (nm)** | 75.46 ± 1.83 | 102.67 ± 1.07 | 107.62 ± 3.47 | 92.18 ± 0.59 | 96.46 ± 1.90 |
| **Tail length (nm)** | 117.42 ± 2.26 | 116.23 ± 1.29 | 123.80 ± 3.96 | 129.14 ± 8.25 | 131.03 ± 1.05 |
| **Phage morphology** | Myovirus | Myovirus | Myovirus | Myovirus | Myovirus |
| ***Bio-Physicochemical properties*** | | | | | |
| **Temperature stability** | 4-55 °C | 4-55 °C | 4-55 °C | 4-70 °C | 4-70 °C |
| **Stable pH range** | 4-10 | 4-10 | 4-11 | 3-11 | 3-9 |
| **50% Adsorption** | 2-4 min | 0-2 min | 6-8 min | 0-2 min | 4-6 min |
| **Latent period** | 20 min | 10 min | 30 min | 20 min | 20 min |
| **Burst Size** | 62.95 ± 17.42 | 915.40 ± 252.22 | 556.02 ± 71.40 | 92.23 ± 51.89 | 414.54 ± 183.23 |
| ***Genomic Properties*** | | | | | |

| | | | | | |
|---|---|---|---|---|---|
| **Genome Size** | 170501 bp | 169720 bp | 356482 bp | 170021 bp | 171101 bp |
| **CDS** | 279 | 271 | 588 | 290 | 283 |
| **GC content** | 39.55% | 39.65% | 35.60% | 40.38% | 40.26% |
| **tRNA** | 4 | 2 | 4 | 2 | 2 |
| **Genome completeness by checkV** | 100% (Complete) | 99.85% (High-Quality) | 100% (Complete) | 99.98% (High-Quality) | 100% (Complete) |
| **Taxonomy predicted by taxMyPhage and comparative studies** | Viruses; Duplodnaviria; Heunggongvirae; Uroviricota; Caudoviricetes; Pantevenvirales; Straboviridae; Tevenvirinae; Dhakavirus | Viruses; Duplodnaviria; Heunggongvirae; Uroviricota; Caudoviricetes; Pantevenvirales; Straboviridae; Tevenvirinae; Dhakavirus | Viruses; Duplodnaviria; Heunggongvirae; Uroviricota; Caudoviricetes; Asteriusvirus | Viruses; Duplodnaviria; Heunggongvirae; Uroviricota; Caudoviricetes; Pantevenvirales; Straboviridae; Tevenvirinae; Gaprivervirus | Viruses; Duplodnaviria; Heunggongvirae; Uroviricota; Caudoviricetes; Pantevenvirales; Straboviridae; Tevenvirinae; Gaprivervirus |
| **Lifecycle predicted by Phage AI** | Virulent (100%) | Virulent (100%) | Virulent (100%) | Virulent (100%) | Virulent (100%) |
| **AMR, Virulence, lysogeny genes** | None | None | None | None | None |
| **Biosample accession** | SAMN45030591 | SAMN47596685 | SAMN45050877 | SAMN45050877 | SAMN47596686 |
| **Genbank ID** | PQ724394 | PV467748 | PV467749 | PQ724392 | PV419677 |
| **Phage with the highest intergenomic similarity (VIRDIC)** | Enterobacteria phage JS10, NC_012741.1 (91.8%) | *Escherichia* phage WG01, KU_878968.1 (92.6%) | *Escherichia* phage PBECO 4, NC_027364.1 (77.5%) | *Escherichia* phage pEC M719 6WT.2, OQ_845958.1 (96.4%) | *Escherichia* phage EAWsc9, PQ_341120.1 (90.2%) |

| | | | | | |
|---|---|---|---|---|---|
| **Similar phage identified by BLASTn with NCBI database (query coverage, %identity)** | *Escherichia* phage dhaeg, MN850609.1 (91%, 96.21%) | *Escherichia* phage vB_EcoM_FB, MT682711 (94%, 96.76%) | NA | *Escherichia* phage ECML-359, OL631482.1 (98%, 98.07%) | *Escherichia* phage ECML-359, OL631482.1 (92%, 97.55%) |
| **Host identified by Phage AI** | *Escherichia coli* | *Escherichia coli* | *Escherichia coli* | *Escherichia coli* | *Escherichia coli* |

**Table S2:** Annotation of all proteins for five isolated *Escherichia* phages.

**Table S3:** Result of VipTree showing similarity between our phages and already existing phages ($S_g$).

**Table S4:** Intergenomic similarity table obtained through VIRDIC.

**Table S5:** VICTOR master table depicting species, genus, and family clusters of the 5 *Escherichia* phage and 49 related genomes, obtained for the D0 formula.

**Table S6:** Sample location from which avian pathogenic *Escherichia coli* (APEC) were isolated and identified bacterial serotypes.

| S.no | Bacterial strain | State/UT | Latitude | Longitude | Serotype |
|---|---|---|---|---|---|
| **1** | bONBS3 | Odisha | 21°48'34.8"N | 84°16'42.3"E | O101 |
| **2** | bONBS1 | Odisha | 21°48'34.8"N | 84°16'42.3"E | O101 |
| **3** | bOVES3 | Odisha | 21°51'23.0"N | 83°59'50.0"E | O11 |
| **4** | bOPPS3 | Odisha | 22°03'18.7"N | 83°49'58.7"E | O119 |
| **5** | bOPPS1 | Odisha | 22°03'18.7"N | 83°49'58.7"E | O119 |
| **6** | bOTLS1 | Odisha | 22°09'17.9"N | 84°02'40.4"E | O9 |
| **7** | bONBS2 | Odisha | 21°48'34.8"N | 84°16'42.3"E | O7 |
| **8** | bOTLS3 | Odisha | 22°09'17.9"N | 84°02'40.4"E | O7 |
| **9** | bDAUS1 | Delhi | 28°30'26.9"N | 77°10'49.9"E | O101 |
| **10** | bDKRS2 | Delhi | 28°39'50.1"N | 77°17'03.9"E | O7 |
| **11** | bBBMV1 | Bihar | 26°37'33.0"N | 84°52'37.4"E | O109 |
| **12** | bBBMV3 | Bihar | 26°37'33.0"N | 84°52'37.4"E | O109 |
| **13** | bBCCV3 | Bihar | 26°38'37.9"N | 84°54'14.5"E | O101 |
| **14** | bBCCV2 | Bihar | 26°38'37.9"N | 84°54'14.5"E | O120 |
| **15** | bBCCV1 | Bihar | 26°38'37.9"N | 84°54'14.5"E | UT |
| **16** | bBZKV1 | Bihar | 26°36'42.1"N | 84°53'44.4"E | O84 |
| **17** | bBBMV2 | Bihar | 26°37'33.0"N | 84°52'37.4"E | O120 |
| **18** | bUMIM3 | Uttar Pradesh | 28°27'33.3"N | 77°56'27.0"E | O101 |
| **19** | bUJBS1 | Uttar Pradesh | 25°30'20.9"N | 82°14'12.8"E | O101 |
| **20** | bUTSS1 | Uttar Pradesh | 25°26'23.6"N | 81°50'38.0"E | O111 |
| **21** | bUJBS2 | Uttar Pradesh | 25°30'20.9"N | 82°14'12.8"E | O169 |
| **22** | bUASS1 | Uttar Pradesh | 25°30'25.9"N | 81°57'20.1"E | O156 |
| **23** | bUAMS3 | Uttar Pradesh | 25°30'46.5"N | 82°13'06.3"E | O128 |
| **24** | bUTSS3 | Uttar Pradesh | 25°26'23.6"N | 81°50'38.0"E | O17 |
| **25** | bUTSS5 | Uttar Pradesh | 25°26'23.6"N | 81°50'38.0"E | O7 |
| **26** | bUMIM1 | Uttar Pradesh | 28°27'33.3"N | 77°56'27.0"E | O20 |

| | | | | | |
|---|---|---|---|---|---|
| **27** | bUASS2 | Uttar Pradesh | 25°30'25.9"N | 81°57'20.1"E | O119 |
| **28** | bPGOV3 | Punjab | 30°09'06.3"N | 74°51'45.0"E | O7 |
| **29** | bPGOV1 | Punjab | 30°09'06.3"N | 74°51'45.0"E | UT |
| **30** | bPGOV2 | Punjab | 30°09'06.3"N | 74°51'45.0"E | O84 |
| **31** | bPGOV4 | Punjab | 30°09'06.3"N | 74°51'45.0"E | O120 |
| **32** | bPMLV1 | Punjab | 30°01'31.7"N | 74°47'53.6"E | O16 |
| **33** | bPMLV2 | Punjab | 30°01'31.7"N | 74°47'53.6"E | O16 |
| **34** | bPMLV3 | Punjab | 30°01'31.7"N | 74°47'53.6"E | O16 |
| **35** | bRAPR3 | Rajasthan | 26°17'53.2"N | 73°03'14.3"E | O120 |
| **36** | bRBJR1s | Rajasthan | 26°16'34.1"N | 72°59'08.7"E | O101 |
| **37** | bRSPR3s | Rajasthan | 26°15'41.8"N | 72°57'42.8"E | O111 |
| **38** | bRSPR1s | Rajasthan | 26°15'41.8"N | 72°57'42.8"E | O109 |
| **39** | bRBJR2s | Rajasthan | 26°16'34.1"N | 72°59'08.7"E | O119 |
| **40** | bRAPR1s | Rajasthan | 26°17'53.2"N | 73°03'14.3"E | O101 |
| **41** | bRAPR3s | Rajasthan | 26°17'53.2"N | 73°03'14.3"E | O101 |
| **42** | bRBKR1 | Rajasthan | 26°16'12.1"N | 73°02'35.7"E | O156 |
| **43** | bRAPR2s | Rajasthan | 26°17'53.2"N | 73°03'14.3"E | UT |
| **44** | bRBJR3s | Rajasthan | 26°16'34.1"N | 72°59'08.7"E | UT |
| **45** | bRBJR4s | Rajasthan | 26°16'34.1"N | 72°59'08.7"E | UT |
| **46** | bRBJR2 | Rajasthan | 26°16'34.1"N | 72°59'08.7"E | O84 |
| **47** | bRSMR1s | Rajasthan | 26°14'28.0"N | 72°57'05.1"E | O5 |
| **48** | bRSMR2s | Rajasthan | 26°14'28.0"N | 72°57'05.1"E | O5 |
| **49** | bRBHR3 | Rajasthan | 26°13'46.5"N | 73°00'08.0"E | O18 |
| **50** | bRAPR2 | Rajasthan | 26°17'53.2"N | 73°03'14.3"E | O169 |
| **51** | bRBJR1 | Rajasthan | 26°16'34.1"N | 72°59'08.7"E | O17 |

**Table S7:** Pearson correlation between the antibiotics and phages activity based on their susceptibility against the 51 APEC isolates.

| | CIP | AMC | NIT | CL | CAZ | COT | TE | ETP | PIT | CXM | fRPOT1 | fBSZT1 | fDMYT1 | fUAMT1 | fPKPT2 |
|---|---|---|---|---|---|---|---|---|---|---|---|---|---|---|---|
| **CIP** | 1.000 | | | | | | | | | | | | | | |
| **AMC** | 0.365 | 1.000 | | | | | | | | | | | | | |
| **NIT** | 0.627 | 0.568 | 1.000 | | | | | | | | | | | | |
| **CL** | 0.499 | 0.340 | 0.422 | 1.000 | | | | | | | | | | | |
| **CAZ** | 0.252 | 0.383 | 0.136 | 0.065 | 1.000 | | | | | | | | | | |
| **COT** | 0.307 | 0.191 | 0.124 | 0.378 | -0.008 | 1.000 | | | | | | | | | |
| **TE** | 0.379 | 0.065 | -0.042 | 0.066 | 0.152 | 0.465 | 1.000 | | | | | | | | |
| **ETP** | 0.139 | 0.268 | 0.213 | 0.210 | 0.305 | 0.164 | 0.049 | 1.000 | | | | | | | |
| **PIT** | 0.288 | 0.614 | 0.451 | 0.477 | 0.296 | 0.227 | 0.073 | 0.565 | 1.000 | | | | | | |
| **CXM** | 0.284 | 0.381 | 0.198 | 0.326 | 0.491 | 0.103 | 0.129 | 0.346 | 0.485 | 1.000 | | | | | |
| **fRPOT1** | 0.228 | -0.097 | 0.023 | -0.056 | 0.267 | 0.056 | 0.372 | 0.010 | 0.097 | 0.136 | 1.000 | | | | |
| **fBSZT1** | 0.291 | -0.065 | 0.042 | 0.020 | 0.188 | 0.168 | 0.425 | 0.150 | 0.175 | 0.188 | 0.889 | 1.000 | | | |
| **fDMYT1** | 0.385 | 0.005 | 0.097 | 0.037 | 0.259 | 0.265 | 0.617 | 0.074 | 0.060 | 0.209 | 0.756 | 0.850 | 1.000 | | |
| **fUAMT1** | 0.270 | -0.023 | 0.089 | -0.005 | 0.383 | 0.101 | 0.460 | -0.029 | 0.039 | 0.223 | 0.846 | 0.803 | 0.817 | 1.000 | |
| **fPKPT2** | 0.323 | -0.097 | 0.103 | 0.030 | 0.013 | 0.056 | 0.294 | 0.010 | 0.014 | 0.057 | 0.529 | 0.652 | 0.756 | 0.610 | 1.000 |

**Table S8:** List of primers used in the study and their sequence.

| S.no | Gene | Primer | Sequence | Annealing temperature | Product Size (bp) |
|---|---|---|---|---|---|
| **1** | ***omp*T** | Forward | TCATCCCGGAAGCCTCCCTCACTACTAT | 60°C | 496 |
| | | Reverse | TAGCGTTTGCTGCACTGGCTTCTGATAC | | |
| **2** | ***iro*N** | Forward | AATCCGGCAAAGAGACGAACCGCCT | 62°C | 553 |
| | | Reverse | GTTCGGGCAACCCCTGCTTTGACTTT | | |
| **3** | ***hly*F** | Forward | GGCCACAGTCGTTTAGGGTGCTTACC | 60°C | 450 |
| | | Reverse | GGCGGTTTAGGCATTCCGATACTCAG | | |
| **4** | ***iut*A** | Forward | GGCTGGACATCATGGGAACTGG | 62°C | 302 |
| | | Reverse | CGTCGGGAACGGGTAGAATCG | | |
| **5** | ***iss*** | Forward | CAGCAACCCGAACCACTTGATG | 58°C | 323 |
| | | Reverse | AGCATTGCCAGAGCGGCAGAA | | |
| **6** | **Bacterial GAPDH** | Forward | TCAGCGTTAATCGCAATCAG | 60°C | NA |
| | | Reverse | GCGACACCCTTTACCTGTGT | | |
